# Regulatory innovation and transcriptome turnover drive the evolution of multicellularity in brown algae

**DOI:** 10.64898/2026.04.27.720984

**Authors:** Jaruwatana Sodai Lotharukpong, Min Zheng, Remy Luthringer, Daniel Liesner, Kenny A. Bogaert, Masakazu Hoshino, Enrique López-Gómez, Noé Fernández-Pozo, Fabian B. Haas, Susana M. Coelho

## Abstract

How complex multicellularity evolved repeatedly across eukaryotes remains a central unresolved question in biology. Whether principles inferred largely from animals and plants reflect universal features of multicellular evolution or lineage-specific outcomes is unknown. Here, using a stage- and tissue-resolved transcriptomic atlas spanning the full life cycles of 15 species across the brown algal radiation, comprising 639 RNA-seq libraries, we uncover general principles governing the evolution of developmental programs in an independently evolved multicellular lineage. Despite broad conservation of genome architecture and synteny, developmental and morphological diversification is accompanied by pervasive rewiring of gene expression and rapid turnover of co-expression networks, identifying regulatory evolution as a major driver of multicellular innovation. We show that evolutionarily young genes, including genes of giant viral origin, are repeatedly recruited into developmental programs, particularly in motile unicellular stages, revealing a broader role for these stages as hotspots of evolutionary novelty. We further identify lineage-specific cis-regulatory innovation linked to active chromatin and large-scale transcriptional rewiring, providing a mechanistic basis for transcriptome turnover. Together, these findings show that brown algae reinvented complex multicellularity through distinct molecular routes shaped by rapid regulatory innovation, while revealing organizational principles conserved across independent multicellular lineages.

## Introduction

The emergence of novelty is a foundational problem in the study of multicellular organisms ^1^. At the genomic level, multi-species comparative analysis of genome content and gene expression patterns have started to shed light on how novelty arises in animals, plants and fungi. For example, *de novo* evolution of coding sequences facilitate the evolution of new traits in animals and plants ^2,3^; reuse of ancient gene families underpins morphological innovations ^4^; joint multi-species analysis of gene repertoire and expression evolution has uncovered systems-level imprints of novelty and constraints in embryo development ^5–7^, organogenesis ^8,9^, and cell and tissue type diversity ^10–12^. However, it is unknown whether common patterns observed in animals and plants represent core principles behind complex multicellular evolution without data from other complex multicellular lineages.

Brown algae (Phaeophyceae) are marine eukaryotes that have evolved complex multicellularity independently from animals, green plants, fungi and red algae ^13,14^, approx. 450 Ma ^15^, providing fertile grounds to explore the generalities of complex life ^7,16^. Since the emergence of complex multicellularity, brown algae have diversified to over 2000 extant species, resulting in a distinct array of morphological forms, sexual systems, life cycle types and number of cell types ^17–19^. Despite this diversification, brown algal genome structures remain conserved, as evidenced by extensive linear genomic synteny and lack of whole genome duplications ^20,21^. Thus, the rapid evolution and frequent turnover of these developmental traits ^17,18^ may have arisen from evolutionary changes to the gene expression landscape, independent of genome structure changes associated with morphological and regulatory evolution in other lineages ^22–25^. Testing this scenario requires RNA-seq data spanning tissues, developmental stages and sexes, which remains scarce outside of a few select species.

Here, we present a comprehensive brown algal RNA-seq dataset, encompassing 15 species that span 450 million years of evolution and capture the full breadth of morphological and sexual diversity within the group. By integrating these new transcriptomic resources with current knowledge of brown algal genome biology, we reveal key patterns and principles underlying transcriptome evolution across this key multicellular lineage. First, we uncover substantial divergence in the expression of shared orthologous genes, across shallow and deep evolutionary timeframes. We demonstrate that novel, taxonomically restricted genes including virus-derived genes are readily incorporated into the tissue and developmental stage-specific transcriptomes, facilitated by their extensive and rapid integration into the gene co-expression network. Furthermore, this co-expression network is fast evolving, with limited conservation of co-expression modules. Sex-related networks evolve particularly fast, as evidenced by the change in network centrality of the master male-determinant factor, *MIN*. Lastly, we detect a step-wise evolution of novel regulatory regions across the 15 species, coinciding with the diversification of the brown algal lineage. Together, our findings show that the remarkable morphological and sexual diversification of brown algae has been driven by rapid transcriptome and regulatory evolution despite broad conservation of genome architecture, revealing an alternative molecular route to complex multicellular diversification.

## Results

### Brown algal transcriptomes evolve rapidly

Brown algae provide a powerful system to study developmental and sexual evolution, including the alternation of multicellular gametophyte and sporophyte generations. However, transcriptomic resources across the brown algal radiation remain limited. To address this gap, we generated transcriptomic data from 15 species spanning the diversity of brown algae, including 14 brown algae (Ectocarpus sp. 7, *Scytosiphon promiscuus, Hapterophycus canaliculatus, Sphaerotrichia firma, Fucus serratus, Fucus distichus, Laminaria digitata, Saccharina latissima, Desmarestia herbacea, Desmarestia dudresnayi, Saccorhiza polyschides, Saccorhiza dermatodea, Sphacelaria rigidula*, and *Dictyota dichotoma*), together with the simple filamentous outgroup *Schizocladia ischiensis* (**Fig. 1a, Fig. S1, Table S1**). Transcriptomic data were obtained by RNA-seq across developmental stages, tissues, and sexes (see **Methods**). We further augmented this dataset by uniformly reprocessing previously published raw RNA-seq datasets using the same computational workflow (see **Methods**). In total, we amassed 3.42 TB of gzip-compressed RNA-seq data comprising 41.0 billion reads across 639 libraries. This comprehensive resource is accessible through an online Seaweed Expression Atlas (http://atlas.algae.team; **Fig. S2**) built using EasyGDB^26^, which currently enables genome browsing, BLAST searches, gene annotation queries, and cross-species expression visualization and comparison.

**Fig 1.**
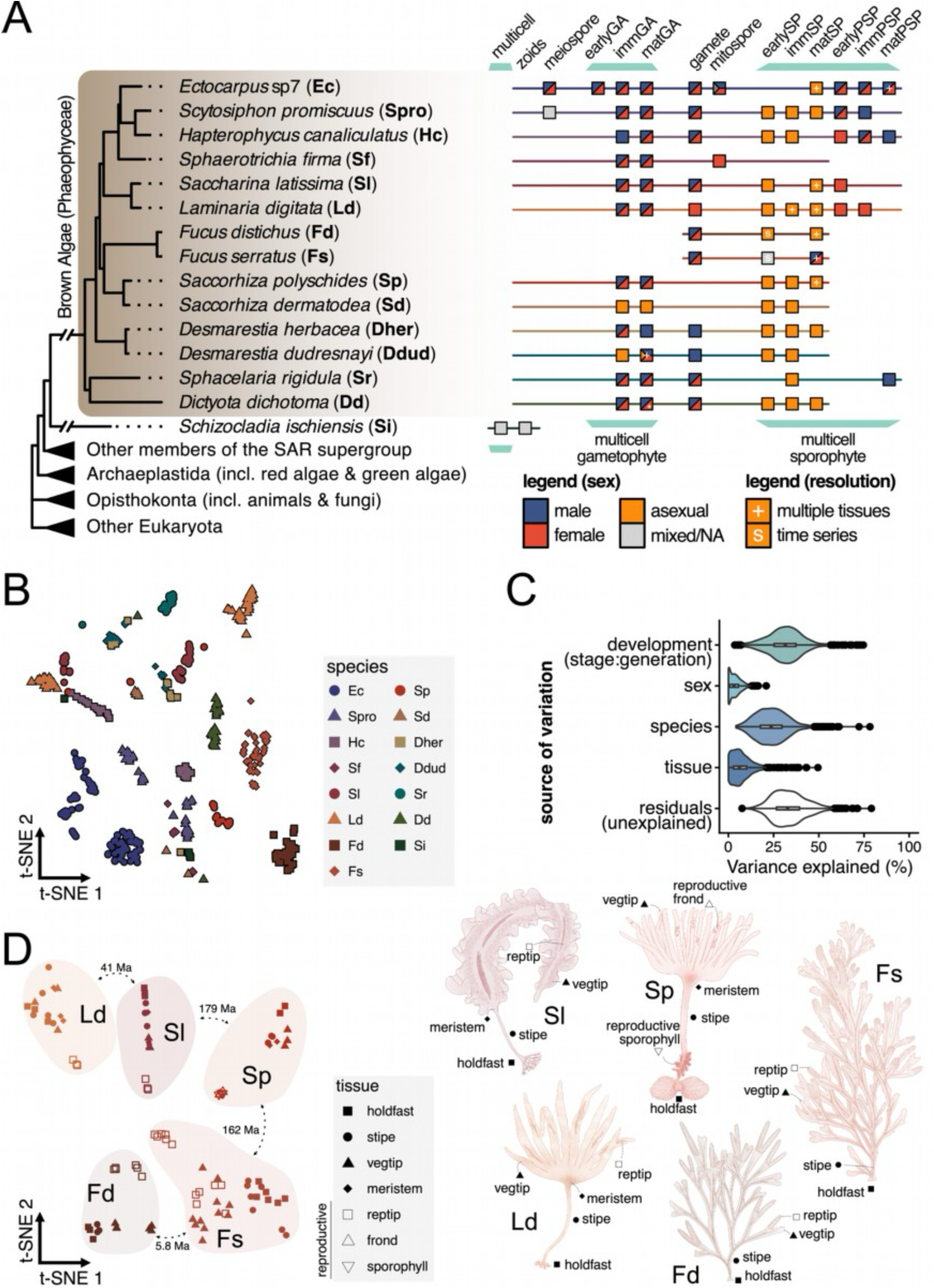
A brown algal gene expression atlas. (**A**) Evolutionary relationship and overview of the transcriptome profiling, across 14 brown algal species and the simple multicellular outgroup, *Schizocladia ischiensis*. These samples span the life cycle, including unicellular and multicellular stages. Gametophytes, sporophytes and parthenosporophytes are indicated by GA, SP and PSP, respectively. Non-gametic zoids (mitospore and meiospore) are coloured based on parent sexual genotype. Mitospore refers to spore derived from plurilocular sporangia while meiospore refers to spore derived from unilocular sporangia. (**B**) A 2D projection (*t*-distributed stochastic neighbour embedding, *t*-SNE) of the RNA-seq dataset across 15 species. (**C**) Variance partition analysis of the RNA-seq dataset. The biological sources of variations considered are development (including generation and developmental stages therein), sex, species and tissue, while the residuals comprise unexplained sources of variation. (**D**) A 2D projection (*t*-SNE) focused on sporophyte tissues across 5 species, accompanied by a visualisation of the different tissues. Reproductive tissues are defined as tissues that produce unicells including meiospores in *L. digitata, S. latissima* and *S. polyschides*) and gametes in (*F. serratus* and *F. distichus*). Each point represents a sample in (**B, D**).

Using this combined dataset, we analyzed transcriptomic relationships across species using 1,455 shared orthogroups. Projection of the multi-species transcriptomic data into a reduced dimensional space revealed that the dominant grouping across the first 20 principal components corresponds to species identity (**Fig. 1b**). Such a strong species signal is indicative of extensive system-wide transcriptomic divergence ^27^. Partitioning the sources of variation using a linear mixed model ^28^ further confirmed that species identity represents one of the strongest contributors to biological variance, alongside developmental stage (**Fig. 1c**). Note that the strength of the species signal was robust to alternative orthology definitions (orthogroups versus 1:1 orthologues) and RNA-seq normalization strategies (**Fig. S3**).

At the same time, the presence of a clear developmental signal raises the possibility that stage-specific transcriptional programs may be partially conserved across species. To explore this possibility, we examined how gene expression varies between life-cycle stages. Most brown algae exhibit complex life cycles that alternate between multicellular haploid gametophyte and diploid sporophyte generations ^18,29^, with considerable variation in the morphology of each phase across species (**Fig. S1**). To examine how life cycle stage contributes to transcriptomic variation, we compared transcriptomes from both generations across species. Within species, expression similarity between generations broadly reflected their degree of morphological resemblance (**Fig. S4a**). Across species, however, gametophytes exhibited higher orthogroup expression correlations than sporophytes (**Fig. S4b, Table S2**). In several cases, sporophyte transcriptomes were more similar to gametophytes of other species than to sporophytes within the same species (**Fig. S4b**). Together, these observations suggest that sporophyte transcriptional programs are less conserved across species, potentially contributing to the greater morphological diversification of sporophytes in brown algae ^18,30^.

To determine whether transcriptomic divergence among sporophytes reflects tissue-specific morphological differences, we compared sporophyte tissues across Laminariales (*L. digitata, S. latissima*), Tilopteridales (*S. polyschides*), and Fucales (*F. serratus, F. distichus*) using 4,188 shared orthogroups. Notably, Tilopteridales and Laminariales represent evolutionarily distinct lineages that have converged on highly similar sporophyte morphologies while occupying comparable ecological niches ^29^, providing a natural system to test whether convergent tissues recruit similar transcriptional programs. Reprojection of the transcriptomes into a two-dimensional space, together with correlation analyses, revealed that samples cluster primarily by species rather than by convergent tissue type (**Fig. 1d, Fig. S4c–d**). In several cases, morphologically dissimilar tissues showed higher expression similarity than convergent tissues (**Fig. S4c–d**). For example, the holdfast of *S. polyschides* (Tilopteridales) displays greater transcriptomic similarity to the stipe and meristem of the same species than to the holdfast of *S. latissima* (Laminariales) (**Fig. S4c**). Thus, morphological convergence between these lineages is not mirrored by convergence in the underlying transcriptome, illustrating that similar morphological structures can arise through distinct gene expression programs.

We next asked whether homologous tissues share transcriptomic profiles across species. Surprisingly, even within Laminariales (*L. digitata* versus *S. latissima*) and within Fucales (*F. serratus* versus *F. distichus*), morphologically similar homologous tissues do not cluster together (**Fig. 1d, Fig. S5a–b**). Within Laminariales, morphologically dissimilar tissues can even show higher expression correlation than homologous tissues across species (**Fig. S5a–b**). These observations indicate that transcriptomic profiles of homologous tissues or developmental stages are largely species-specific, even when morphology, sexual identity, or developmental stage is conserved. Furthermore, comparisons with mammal and flowering plant datasets (**Fig S5c**) ^31,32^ show that transcriptome divergence among brown algal tissues is comparable to that observed in rapidly evolving flowering plants and substantially greater than that seen among mammalian tissues. This pattern supports a rapid rate of transcriptome evolution in brown algal tissues, which can explain the transcriptomic divergence among homologous tissues across shallow evolutionary timescales within Laminariales and Fucales.

Interestingly, across all projections, the sporophyte tissue types showing the lowest divergence correspond to reproductive structures involved in unicell production (**Fig. 1d, Fig. S4c–d, Fig. S5a–b**), displaying more similar expression of orthologous genes than other tissue types. In contrast, somatic sporophyte tissues exhibit substantially greater divergence, consistent with extensive lineage-specific rewiring of gene expression. These observations suggest that functional constraints associated with unicell production, rather than morphological similarity, are more likely to shape transcriptome conservation.

In summary, our analyses reveal a strong species-level signal across both deep and shallow evolutionary timescales in brown algae, with both convergent and homologous somatic tissues recruiting distinct gene sets to achieve similar developmental outcomes, while reproductive tissues involved in unicell production reveal a comparatively conserved transcriptional profile.

### Evolutionary young genes are dynamically expressed

Across the tree of life, novel (taxonomically restricted) genes have been tied to the emergence of new biological traits ^2,3^. In brown algae, a burst of new gene families has been reported, coinciding with the origin of complex multicellularity in this group ^19^, with 29.6% of the gene repertoire of brown algae being unique to this group. In addition, novel genes continuously emerge along brown algal evolution ^20^. However, how these genes are expressed and how they contribute to species-specific transcriptomes remain unclear.

To address this, we analyzed the expression patterns of genes of different evolutionary ages (**Fig. S6**) across developmental stages, tissues, and sexes in brown algae. This analysis revealed a consistent pattern across all species: evolutionarily young genes were disproportionally expressed in a tissue and/or stage specific manner, as quantified using *tau* ^33,34^, compared to older genes in all species (**Fig 2a**). It should be noted that the changes in expression-specificity patterns across gene ages appears gradual, with novel genes becoming increasingly more broadly expressed over evolutionary time. In line with this, the burst of new genes coinciding with the origin of complex multicellularity is not associated with higher expression specificity (**Fig 2a, Fig S6**, see phylorank 8 /Phaeophyceae-level genes). The preferentially specific expression of young genes persisted after controlling for the correlation between mean expression and *tau* (**Fig. S7a–b**), and when restricting the analysis to a subset of young genes that pass the homology detection failure test ^35^ (**Fig. S7c**). Similar expression patterns of novel genes have been reported in animals and plants ^36,37^ suggesting a common trajectory for the functionalization of novel genes across complex multicellular lineages. In this scenario, relaxation of evolutionary and pleiotropic constraints in young genes, initially expressed in a highly specific manner, may facilitate exploration of new molecular and phenotypic space ^2,38^.

**Fig 2.**
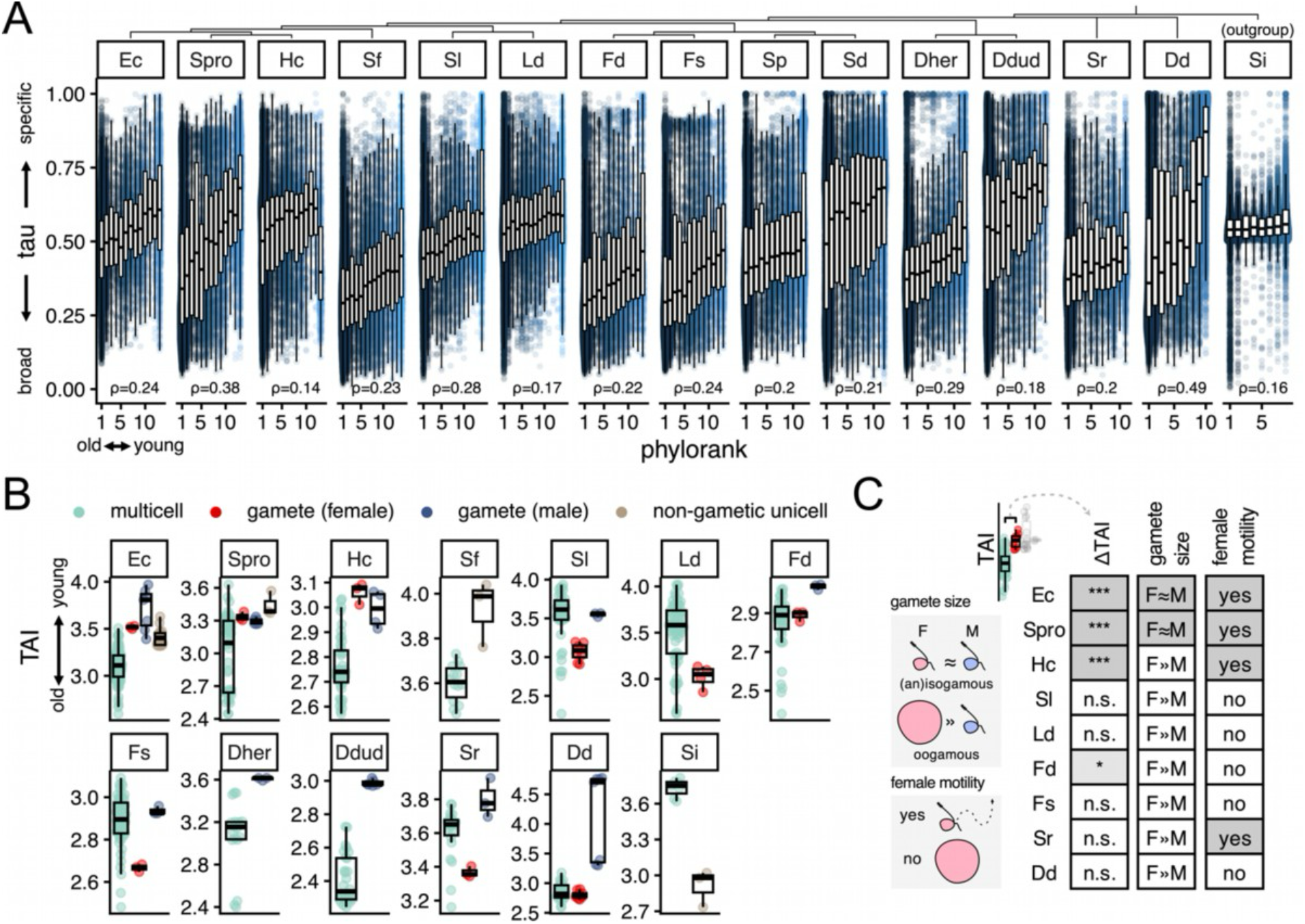
Transcriptomic incorporation of young genes via expression specificity. (**A**) The relationship between expression specificity (*tau*) and gene age (phylorank). Each point represents a gene, with younger genes coloured by lighter shades of blue. Spearman correlation coefficient is indicated using ρ. High *tau* value indicates specific expression (*e*.*g*., stage, sex and/or tissue), while broadly expressed genes are annotated with a low *tau* value. High phylorank indicate evolutionarily young genes, and vice versa for low phylorank. The evolutionary relationship between the species is displayed in the dendrogram above. (**B**) Transcriptome age index (TAI) profiles across the species with both unicellular and multicellular stages. Samples enriched in evolutionarily younger genes have a higher TAI than samples enriched in older genes. Unicellular samples are separated by sex and function (male gametes, female gametes and non-gametic unicellular zoids). (**C**) TAI differences between female gametes and multicellular stages are linked to cell morphology and behaviour. The gamete size difference and mobility are illustrated.

We next investigated where evolutionarily young genes are expressed across developmental stages and tissues by computing the transcriptome age index (TAI), which captures the expression-weighted mean gene age ^5^. Across the 14 brown algal species, TAI values tended to be higher in unicellular samples than in multicellular stages (**Fig. 2b**), indicating that unicellular stages preferentially express younger genes. However, female gametes of *S. latissima, L. digitata, F. serratus, S. rigidula*, and *D. dudresnayi* do not show elevated expression of young genes relative to the average multicellular stage (**Fig. 2c**). Because these species produce large, immotile eggs, this pattern suggests that the enrichment of young genes is associated with gamete motility and relative cell size rather than with unicellularity *per se* (**Fig. 2c**). Consistent with this interpretation, non-gametic zoids of *Ectocarpus, S. promiscuus*, and *S. firma* also show elevated expression of young genes and display phenotypic similarities to motile gametes. It should be noted that TAI is lower in motile unicells compared to (simple) multicellular stages in the outgroup *S. ischiensis* (**Fig 2b**). This suggests that the pattern of young genes in motile unicellular stages applies only to complex multicellular lineages.

In contrast to unicellular stages, multicellular stages show no consistent differences in TAI between gametophytes and sporophytes or between males and females (**Fig. S8a–b**). This absence of a TAI signal is unexpected given the pronounced divergence in orthologue expression observed in sporophytes (**Fig. S4b**), suggesting that the morphological divergence in multicellular stages is driven predominantly by older genes rather than lineage-specific young genes. Together with previous work in other multicellular systems ^39–42^, our results suggest that evolutionarily young genes are consistently enriched in motile unicellular stages, linking their expression to dispersal and reproductive functions across complex multicellular lineages.

### Extensive rewiring of gene co-expression networks

Global transcriptome dynamics are shaped by gene co-regulation, which can be captured using gene co-expression networks. Such networks can reveal the mechanisms underlying transcriptome evolution. For each species with more than 30 libraries, we computed a consensus co-expression network of 12 different methods and algorithms ^43^. We then explored the resulting co-expression network by examining the network centrality of each gene (**Table S3**). We found that functional information is, in part, encoded in the network centrality. For example, across brown algae, TALE homeodomain transcription factors retained high centrality values (**Fig S9a-b**), including experimentally characterised life cycle master regulators in *Ectocarpus, ORO* and *SAM* ^44^. Intriguingly, the centrality of *MIN* orthologues (an HMG-domain master male sex inducer), inferred via reciprocal best hit, corresponded with function ^45^; *MIN* orthologues were central in species with a MIN-determined UV sexual system, whereas the centrality values dropped with the loss of this sexual system ^20^ (**Fig S9c**). Thus, the switch in sexual systems in brown algae is underscored by a rapid evolution of gene networks.

We next asked how evolutionarily young genes are incorporated into the co-expression network by examining the relationship between gene age and network centrality. Strikingly, the range of centrality values do not differ between evolutionarily young and old genes (**Fig 3a**), suggesting a rapid co-option of young genes into the broader transcriptomic network, a pattern reminiscent of observations in unicellular yeasts ^46^. By partitioning the network into information-theoretic communities (or modules) of co-expressed genes ^47^, we found a non-random distribution of gene age by modules (**Fig 3b**). In other words, evolutionarily young genes tend to co-express with each other, resulting in old and young ‘neighbourhoods’ in the co-expression space (**Fig 3c**). Notably, at the module level, young genes were generally more peripheral, except in the module with the youngest genes, wherein the young genes are more central than older genes (**Fig S10a-b**). The presence of old and young modules, explains to some extent why young genes may be central in the overall co-expression network. Together, these results suggest that evolutionarily young genes rapidly integrate into the transcriptome within recently-evolved modules, allowing them to assume central roles while minimising disruption to global network organisation.

**Fig 3.**
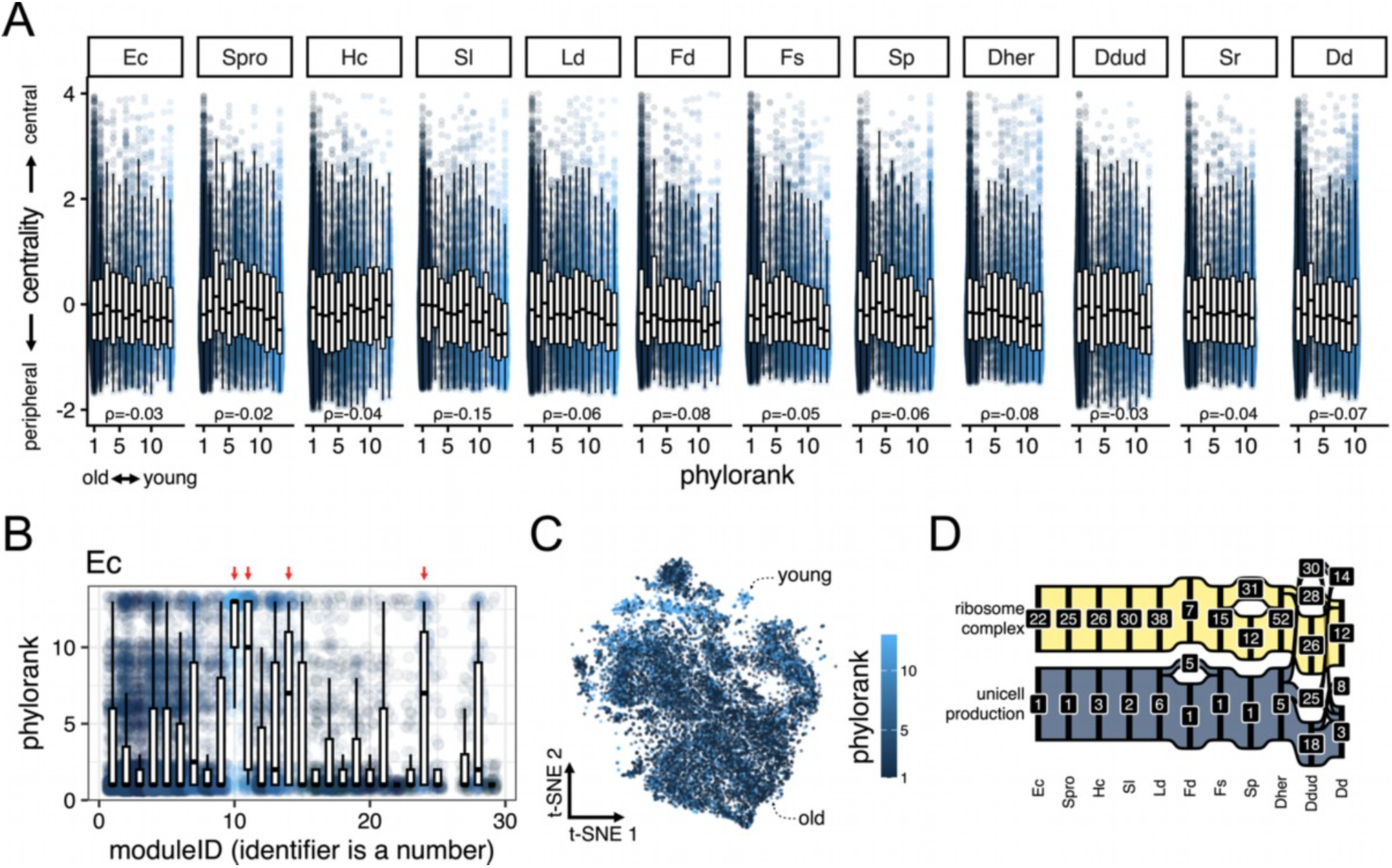
Cross-species comparison of gene co-expression patterns. (**A**) The relationship between gene co-expression network centrality (using z-score of PageRank) and gene age (phylorank). Each point represents a gene, with younger genes coloured by lighter shades of blue. Spearman correlation coefficient is indicated using ρ. Genes at the periphery in the network have low network centrality values, and vice versa for genes central to the network. High phylorank indicate evolutionarily young genes, and vice versa for low phylorank. (**B**) Distribution of gene age (phylorank) across inferred gene co-expression modules in *Ectocarpus*. Module names are given as number. Red arrows highlight young modules (median gene age is greater than five). Each point, representing a gene, is coloured by the mean phylorank of the module. (**C**) Distribution of gene age (phylorank) across the 2D projection (*t*-SNE) of genes in *Ectocarpus* using expression data. Each point represents a gene, with younger genes coloured by lighter shades of blue. Young and old regions in the projection are labelled for visual clarity. (**D**) Conservation of *Ectocarpus* module 1 and 22 across brown algae, based on overlap of orthologues grouped in the same module. Since module names are given as number, the exact number is unimportant as it varies between species.

In light of several lines of evidence consistent with a fast-evolving transcriptome, we asked if we can still detect evolutionary conservation in the gene network space. To this end, we annotated the co-expression modules through enrichment analysis of protein domains and gene ontology (GO) terms, focusing on the model brown alga *Ectocarpus* (hypergeometric tests; **Table S4**), and examined the evolutionary stability of the co-expression modules. By tracing the co-expression modules of 3500 *Ectocarpus* orthologues across approx. 230 Ma of brown algal evolution (inferred via reciprocal best hit, μ = 8.2k, σ = 1.3k), we detected limited conservation for most modules (**Fig S10c**). However, there were notable exceptions of conserved ancient co-expression modules (**Fig 3d**). *Ectocarpus* module 1 is shared across all species and comprises genes linked to unicell production, as evidenced by enrichment in GO terms associated with cilium assembly (**Table S4**). This result corroborates the deep conservation of this gene set across unicellular and multicellular eukaryotes ^48,49^. It should be noted that genes from this module are not highly expressed in the motile unicell itself but appear in the reproductive structures that produce the unicells (**Table S5**). This pattern supports a scenario where highly conserved genes are expressed in the reproductive structures prior to unicell release, whereas in free-swimming post-release unicells, evolutionarily young genes are enriched, as detected in the TAI analysis (**Fig 2b-c**). Unsurprisingly, this signal of co-expression conservation is also apparent in *Ectocarpus* module 22, which comprise constituents of the ribosome complex (**Table S4**). Thus, despite signals for rapid transcriptome evolution, we were also able to detect deeply conserved co-expression modules linked to ancestral traits of heterokont eukaryotes.

### Horizontally transferred genes integrate into developmental networks

In conjunction with novel, taxonomically restricted genes, horizontal gene transfer (HGT)-derived genes can supply an additional source of novelty in brown algal transcriptomes ^19^. HGT events are widespread in eukaryotes ^50^, contributing to brown algal genome evolution ^19^. Despite the genomic evidence for the retention of HGT-derived genes, the transcriptomic co-option of these HGT-derived genes is uncharacterised in brown algae. We thus examined the expression profile of inferred HGT-derived genes ^19^, focusing on the model brown alga, *Ectocarpus*.

We found that a vast majority of HGT-derived genes is expressed in *Ectocarpus*, with higher proportions for genes stemming from older HGT events (**Fig 4a**). Older HGT-derived genes in *Ectocarpus* also display similar levels of expression specificity to non-HGT genes, while young HGT-derived genes were significantly restricted in expression (**Fig 4b**). Closer inspection of the gene co-expression network revealed that the network centrality of most HGT-derived genes is indistinguishable from other genes in the genome, though young HGT-derived genes were slightly more peripheral (**Fig 4c**). Thus, HGT-derived genes are well-integrated into the gene network. Among the HGT-derived genes with high centrality values (z-score > 2) are genes encoding domains such as Alpha/beta hydrolase, Cyclophilin-like and 4’-phosphopantetheinyl transferase, as well as a more recent HGT-derived gene without domain annotation (**Fig 4d**). Moreover, the co-expression network module with the highest mean centrality of HGT-derived genes is module 5 (**Fig 4d**), which also contains experimentally validated developmental regulators *ORO, SAM* and *IMM* ^44,51^ (**Fig S9d**). This points to a possible role for HGT-derived genes in developmental pathways. Together, these patterns show that HGT-derived genes can persist in brown algal genomes through relatively rapid integration into the host gene regulatory network, initially exhibiting tissue- or stage-specific expression and later expanding their expression breadth.

**Fig 4.**
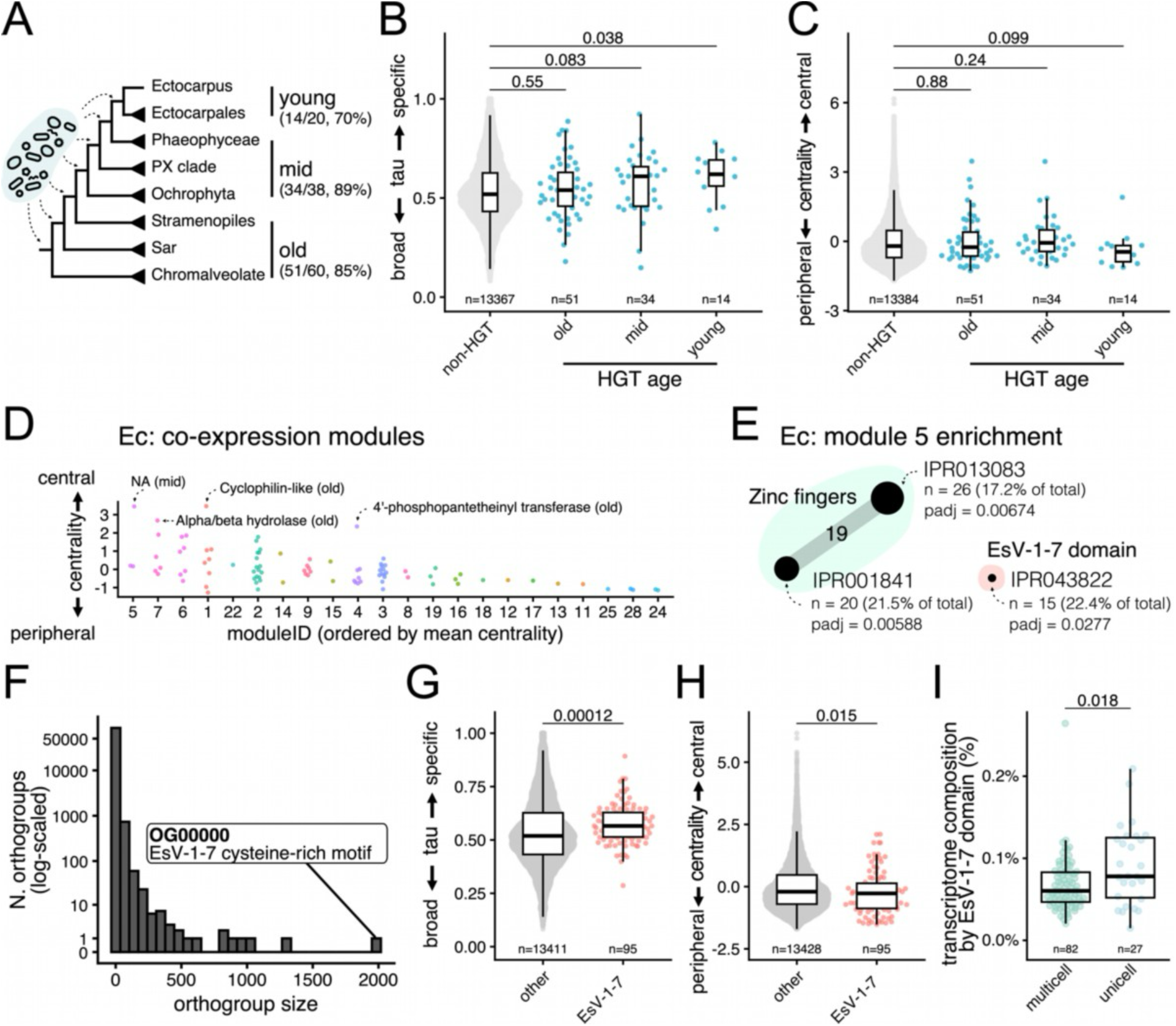
Expression dynamics of HGT-derived genes in brown algae. (**A**) Number of expressed (TPM > 2) HGT-derived genes in *Ectocarpus*, out of total, across evolutionary strata (young = *Ectocarpus* to Ectocarpales, mid = Phaeophyceae to Ochrophyta, old = Stramenopiles to Chromalveolate). Expressed genes are given on the left-hand-side of the bracket; total genes are given on the right-hand-side of the bracket. HGT-derived genes were obtained from ^19^. (**B**) Expression specificity (*tau*) between non-HGT genes and HGT-derived genes of different ages. (**C**) Co-expression network centrality between non-HGT genes and HGT-derived genes of different ages. (**D**) Co-expression network centrality of HGT-derived genes, grouped by module ID. The mean centrality of HGT-derived genes is used to order the module IDs. Genes with high centrality (z-score > 2) are annotated by protein domain information and evolutionary stratum. (**E**) Enriched protein domains in module 5 of the co-expression network in *Ectocarpus*, presented as a co-occurence network. The nodes represent the enriched domain identifier (from InterPro) and are labelled with the number of genes represented by the given domain (“n”), the percentage of all genes with that given domain in the module, and adjusted p-values (“padj”) obtained via hypergeometric tests and the Benjamini-Hochberg procedure. Short descriptors “Zinc Fingers” and “EsV-1-7 domain” summarise the given entries. The edges are labelled with the number of co-occurence, *i*.*e*. gene models that share the domain. (**F**) Histogram of orthogroup size, inferred via orthoHMM. The largest orthogroups with putative virus-derived domains, EsV-1-7 and FNIP, are annotated. (**G**) Expression specificity (*tau*) between genes with EsV-1-7 domain, FNIP domain and background. (**H**) Co-expression network centrality between genes with EsV-1-7 domain, FNIP domain and background. Genes without EsV-1-7 and FNIP domains are grouped as “background” in (**G**-**H**). (**I**) EsV-1-7 domain genes in the transcriptome of each sample, represented as percentage of total TPM belonging to EsV-1-7 domain genes, are grouped by multicellular and unicellular stages. Higher values represent a transcriptome enriched in EsV-1-7 domain genes. The highest % for multicell is a unilocular sporangia sample (that produces unicells, meiospore). *P*-values from two-sided unpaired Wilcoxon tests are given in (**B**-**C, G**-**I**). Each point represents a gene in (**B**-**D, G**-**H**). The total number of genes belonging to each group is labelled (“n”) in (**B**-**C, G**-**H**). The total number of samples belonging to each group is labelled (“n”) in (**I**). Centrality value is defined as the z-score of PageRank over all expressed *Ectocarpus* genes. *Ectocarpus* data is used for (**A**-**C, G**-**I**). Each point represents a sample in (**I**).

Viruses, particularly *Nucleocytoviricota*, are a significant source of HGT in aquatic eukaryotes lacking strict germline segregation ^52^. In brown algae, widespread integration of *Nucleocytoviricota* has been reported across multiple genomes ^19,53^, and is thought to shape their genome evolution through the horizontal transfer of protein domains, including the EsV-1-7 cysteine-rich domain. This domain is present in the *Ectocarpus* developmental regulator *IMM* ^51^ and has been implicated in the evolution of complex multicellularity in brown algae ^54^. Notably, while EsV-1-7 domain-containing genes are outside of the HGT gene set provided in Denoeud et al. (2024), we observe strong support for their viral association via pairwise sequence alignment analysis (**Fig S11a-c**). Given these observations, we investigated whether viral-derived genes contribute to multicellular development in brown algae focusing on EsV-1-7 domain-containing genes and examining the gene co-expression module 5 in *Ectocarpus*, which is associated with known developmental regulators. This analysis revealed a significant enrichment of EsV-1-7 domain genes in this module (**Fig 4e**), consistent with a role in developmental processes. Notably, the largest orthogroup (OG00000) across brown algal genomes is composed of EsV-1-7 domain-containing genes (**Fig 4f**), including the developmental regulator *IMM* (**Fig S11d**), highlighting the functional importance and expansion of this virus-derived domain.

Given the unique expansion of EsV-1-7 domain genes in brown algal genomes, we next asked whether their expression patterns differ from other host genes. In *Ectocarpus*, EsV-1-7 domain-containing genes exhibited higher expression specificity (tau) than the background gene set (**Fig 4g**), were generally more peripheral in the co-expression network (**Fig 4h**) and were preferentially expressed in unicellular stages (**Fig 4i**). This behaviour mirrors that of other evolutionarily young host-origin genes. Notably, two EsV-1-7 genes (*Ec-07_001800* and *Ec-14_004340*) showed particularly high centrality (z-score > 2) and were assigned to the developmental co-expression module (module 5) (**Table S3**). These results give evidence to a potentially important role of viral-derived genes in facilitating multicellular development in brown algae. Overall, the similar expression behaviour of viral HGT-derived EsV-1-7 genes to other novel, taxonomically restricted genes is indicative of functional co-option of putative virus-transferred genes that uniquely characterise the brown algal genome.

### Step-wise evolution of regulatory regions

Lastly, we asked if the fast evolution of gene expression patterns is linked to specific features of the brown algal regulatory genome, such as promoter sequences. By exploring the genomic regions 500bp upstream of the CDS and comparing the 3-letter nucleotide counts with other model eukaryotes (*A. thaliana, C. elegans, D. melanogaster* and *S. cerevisiae*), we noted that *Ectocarpus* occupies a distinct sequence space, while the other eukaryotes cluster together (**Fig 5a**). This separation between *Ectocarpus* and other eukaryotes is also observed when restricting the analysis to 200bp upstream sequences (**Fig S12a**), and is largely lost when analysing 200bp downstream sequences (**Fig S12b**). Moreover, the 14 brown algal species and the outgroup *Schizocladia* occupy similar sequence space (**Fig S12c**). These results indicate that the genomic regions directly upstream of genes in brown algae differ in sequence composition to other model eukaryotes.

**Fig 5.**
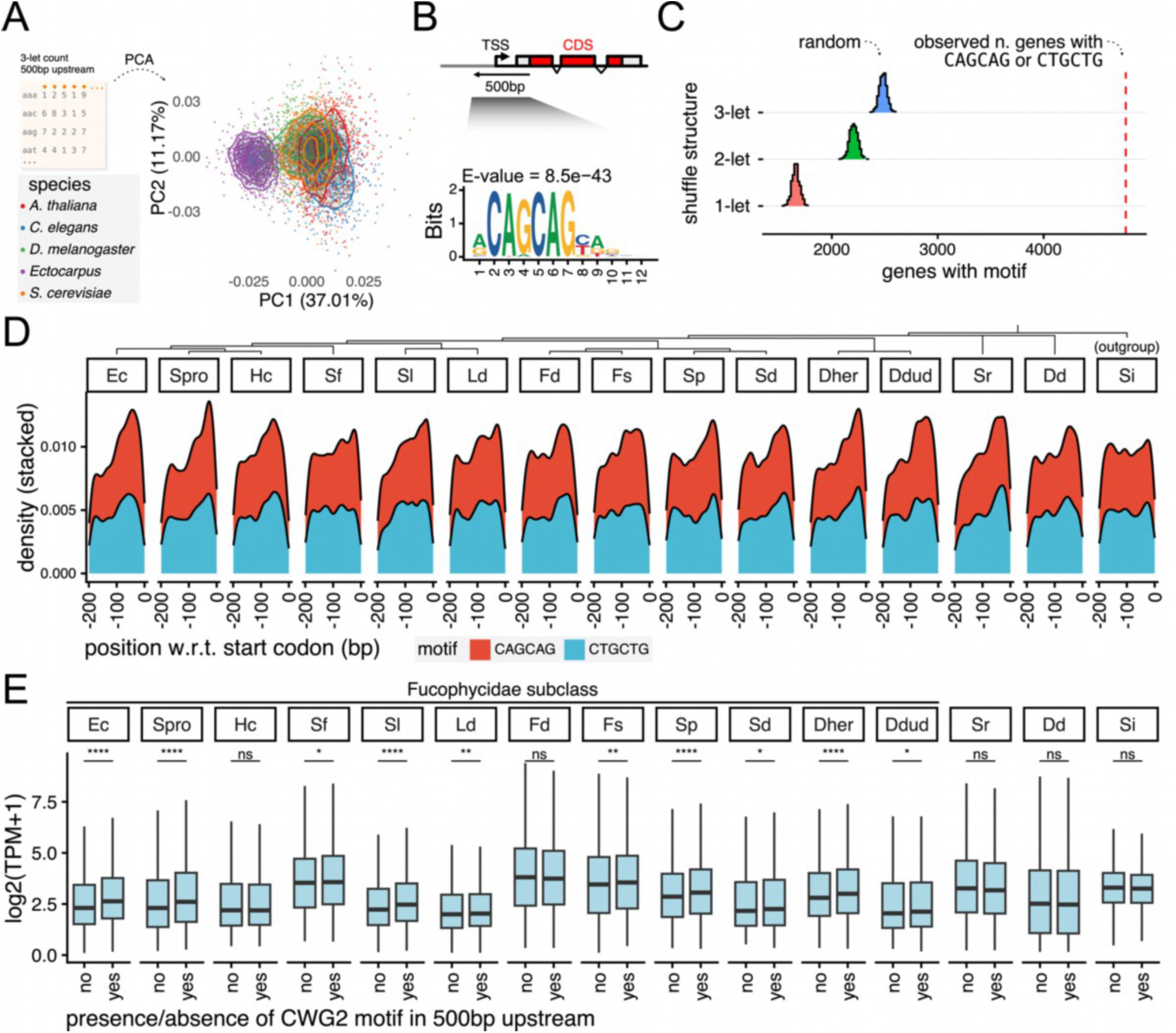
Step-wise acquisition of a distinct regulatory region architecture in brown algae. (**A**) PCA of 3-let sequence composition 500bp upstream of the CDS across *Ectocarpus* and model eukaryotes (*A. thaliana, C. elegans, D. melanogaster* and *S. cerevisiae*). Each dot represents a gene. For visualisation, the number of genes displayed has been subsampled to 500 for each species. (**B**) Enrichment and sequence logo of CAGCAG motif, which partly differentiates *Ectocarpus* from other eukaryotes. (**C**) The distribution of observed CAGCAG (as well as reverse complement CTGCTG) in *Ectocarpus* compared to reshuffled 500bp upstream region using different sequencing re-shuffling strategies (that is, 1-, 2- or 3-letter shuffling strategies). These strategies retain mono-, di- and tri-nucleotide local sequence composition, respectively. The observed motif count is indicated in a red dotted line. (**D**) Stacked density plot of observed positions of CAGCAG and CTGCTG upstream of the CDS across 14 brown algae and outgroup *Schizocladia*. The evolutionary relationship between the species is displayed in the dendrogram above. (**E**) Expression levels (as log2(TPM+1)) of genes with and without CTGCTG or CAGCAG. Members of the Fucophycidae subclass (containing here *Ectocarpus, S. promiscuus, H. canaliculatus, S. firma, S. latissima, L. digitata, F. distichus, F. serratus, S. polyschides, S. dermatodea, D. herbacea* and *D. dudresnayi*) is indicated above. *P*-values were taken from the one-sided Wilcoxon rank sum test; “ns”, not significant, *, p < 0.05; **, p < 0.01; ***, p < 0.001; ****, p < 0.0001. Gene models that overlap within +/-1kb were excluded from (**C**-**E**).

Consistent with the distinct sequence composition of brown algal regulatory region, we found a lack of enrichment for canonical motifs such as the TATA-box *sensu stricto* (**Fig S12d**). This may relate to previous challenges in promoter identification in *Ectocarpus* ^14^. In place of these expected motifs, a previously unreported CWGCWG motif (W = A or T; that is, either CAGCAG or CTGCTG) was detected (**Fig 5b, Fig S12e**), present 500bp upstream of the CDS in 71.3% of genes in the model alga *Ectocarpus* (**Fig S12e**). The CDS was chosen to reduce the impact of 5’ UTR annotation quality differences between brown algal species. The reported motif has no exact match with other known motif, though, a motif similarity search revealed C2H2 zinc finger factor binding sites (MA2532.1, MA1628.2 and MA0697.3) as the closest matches (see **Methods**). The enrichment of CWGCWG motif in the 500bp upstream region is further supported by comparing observed motif count against empirical null distributions obtained by re-shuffling each 500bp region while preserving mono-, di-, or trinucleotide composition ^55^ (**Fig 5c**). Positionally, the CWGCWG motif is commonly detected within 100bp upstream of the CDS (**Fig 5d, Fig S13a**), except in the outgroup *Schizocladia*. We also note a secondary accumulation of the motif at the transcription end site in *Ectocarpus* (**Fig S13a**). In line with the upstream positional enrichment, orthogonal analysis of the overlap between CWGCWG motif sites across the genome and ChIP-seq data in *Ectocarpus* indicates that the CWGCWG motif is associated within H3K4me3 and H3K9ac chromatin contexts (**Fig S13b-c**), particularly among genes with an overall “active” chromatin signature (**Fig S13d**). Furthermore, the motif was depleted of H3K79me2 (**Fig S13b-c**), which is a repression-associated mark in brown algae ^56–58^ that marks intact transposable elements ^59^. This orthogonal analysis raises the possibility that the CWGCWG motif plays a role in chromatin modification pathways.

Given the association of the CWGCWG motif with transcriptionally active chromatin marks in *Ectocarpus*, we examined whether the presence of this motif impacts average gene expression levels across brown algae. Despite the similar positional profile across all brown algae (**Fig 5d**), the presence of this motif correlates with expression levels only within the Fucophycidae subclass (which includes *Ectocarpus, S. promiscuus, H. canaliculatus, S. firma, S. latissima, L. digitata, F. distichus, F. serratus, S. polyschides, S. dermatodea, D. herbacea* and *D. dudresnayi*), with the exception of *H. canaliculatus* and *F. distichus* (**Fig 5e**). This pattern suggests a stepwise evolutionary process: the CWGCWG motif originated in brown algae, later followed by association with increased transcript level in the Fucophycidae subclass. Overall, these results highlight the variation in expression-associated sequence determinants across eukaryotes. Alongside recent work on the distinct epigenetic controls of gene expression in brown algae ^58^, our data posits that the brown algal radiation is characterised by an altered regulatory environment that drives gene expression patterns and, with it, their evolution.

## Discussion

Here, we characterised the transcriptome and its evolution in brown algae, a distinct eukaryotic radiation with evolutionary and ecological significance ^19^. We detected extensive transcriptomic turnover when comparing the expression and co-expression of shared orthologous genes. Moreover, evolutionarily young genes (not shared between species), as well as genes of putative viral origin, are readily incorporated into the transcriptome through tissue or stage specific expression. These patterns are independent of sexual system or levels of morphological complexity, and may be supported by the evolving landscape of regulatory regions in this lineage. Overall, our data supports a scenario where the rapid evolution of organismal traits in brown algae is deeply tied to a rapidly evolving transcriptome, with minimal influence from the slow evolution of their underlying genome structure ^20,21^.

All datasets generated in this study including gene expression, orthology inference, co-expression network metrics, gene age, functional annotations, and upstream genomic sequences are available through the Seaweed Expression Atlas http://atlas.algae.team. Together, this resource and our analysis of brown algal transcriptome evolution provide a valuable community resource for future studies of developmental processes ^60^, host–virus interactions ^61^, and cell type diversity in complex multicellular brown algae ^62^.

A key finding of this study is the distinctive expression pattern of evolutionarily young genes. Recent analyses of brown algal genomes have linked the emergence of complex multicellularity in this lineage to a burst of new gene families ^19^, mirroring patterns observed during multicellular transitions in other lineages ^63–65^. Our data extends these findings by providing transcriptional evidence that these evolutionarily young genes are actively deployed across developmental stages, tissue types and sexes. Moreover, we identify a consistent evolutionary pattern: young genes, and particularly horizontally transferred and virus-derived genes, tend to exhibit more restricted expression across tissues, developmental stages, and sexes than older genes. Importantly, this pattern persists after controlling for the correlation between mean expression and expression specificity, indicating that it cannot be explained solely by ‘leaky’ transcription, i.e, low-level RNA production resulting from non-selective transcriptional activity.

We further identify motile unicellular stages as the predominant context in which both young genes and virus-derived genes are expressed. In animals and plants, young genes have previously been found to be preferentially expressed in male gametes ^8,39– 41^. However, in these systems it is difficult to disentangle the factors underlying this pattern, as male gametes are simultaneously male, motile, and small. Brown algae provide a natural system in which these traits are partially decoupled. In addition to the canonical combination of small motile male gametes and large immotile female gametes, several species produce motile female gametes and/or non-gametic zoids.

Leveraging this biological diversity, our data show that the expression bias of young genes extends beyond sperm to all motile unicellular stages, irrespective of sex. This observation suggests that the “out-of-testis” hypothesis for the initial expression of new genes in male reproductive cells ^66,67^ may partly reflect the size and functional properties of motile cells rather than sex alone. Taken together, our findings suggest that motile unicellular stages act as a “nursery” for evolutionarily young genes, some of which may subsequently provide the raw genetic material for developmental innovation and diversification in brown algae.

The functionalisation process of young genes is also reflected in their rapid integration into the co-expression network, as evidenced by the similar range of centrality values displayed by both young and old genes. We found that these young genes were typically contained within a small subset of gene network modules. This modular organisation in the co-expression space of young genes has been observed in other systems such as the nematode *Pristionchus pacificus*, and is thought to reflect a mechanism for the co-option of orphan genes ^68^. In other words, this modular network organisation may allow young genes to evolve in expression patterns without disrupting the global transcriptome. Together with the specific expression patterns of young genes, these observations may represent broader principles of modularity and evolvability in transcriptome evolution patterns across complex multicellular eukaryotes.

Since the relatively recent diversification of brown algae compared to other complex multicellular lineages ^15^, the brown algae lineage is characterised by frequent turnover of organismal traits such as sexual system, sexual dimorphism, life cycle types and morphological complexity ^17–19^. These rapid changes are not surprising in light of our observation of rapid transcriptome evolution in this lineage, corroborated by previous findings in a more limited set of brown algal species ^30,69,70^. Mechanistically, these changes may be tied to the transcriptomic integration process of young genes, whose incorporation could drive interspecies transcriptomic divergence of shared orthologues in brown algae through concerted evolution ^27^. Our genomic analysis further implicates the role of a shifting regulatory landscape as a contributor to transcriptomic divergence. Importantly, we detected a distinct sequence space 5’ upstream of genes in brown algae, enriched in a novel CWGCWG motif. This motif is over-represented within H3K4me3 and H3K9ac contexts, and is associated with increased expression levels within the broader Fucophycidae subclass (which includes most of the brown algal species in this study). This finding points to a step-wise evolutionary incorporation of a new genomic signal, though the molecular role of this sequence remains unknown.

Together, our data supports the view that changes in the mode of regulation, also recapitulated in the changes to epigenetic controls in brown algae ^58^, can rewire gene regulation for all (including ancient) genes, contributing to a transcriptome evolution pattern distinct to this lineage.

Moreover, organism-level traits may also shape transcriptomic divergence. For example, the rapid transcriptome evolution in land plants is thought to be linked to organismal traits associated with sessile organisms that constantly interact with the external environment ^32^. These features also characterise brown algal biology. Brown algae display remarkable developmental plasticity ^60,71^, where organismal forms are modulated by mechanical forces. This suggests that the transcriptome may present a baseline for morphological potentials, later entrained by environmental cues ^72,73^. In this view, environmental feedback on the gene regulatory circuits may contribute to morphological convergence between sporophytes of Tilopteriales and Laminariales. This developmental flexibility also extends to a lack of a strict germline barrier in brown algae (*i*.*e*., the gamete-producing cells do not appear predetermined nor ‘protected’ in early development), which may relax the control against genomic changes that incorporates novel and HGT-derived genes.

However, the patterns of transcriptomic divergence are not distributed evenly across different tissues or gene classes. In sporophytes, gene expression patterns in somatic tissues evolve faster than unicell-producing reproductive tissues, indicating that functional constraints of producing unicells can shape transcriptome divergence more than morphological resemblance. This finding suggests that somatic tissues are freer to explore new molecular space, potentially facilitating ecological and morphological diversification. Furthermore, despite the general pattern of transcriptomic divergence, analysis of co-expression networks revealed evolutionary persistence of network modules involved in unicell production, which can contribute to the relative transcriptomic conservation of reproductive tissues. Such cellular processes are ancestral to heterokont eukaryotes, underscoring the balance between deep evolutionary constraints and innovations in both gene repertoire and expression in shaping multicellular traits in this independent eukaryotic radiation.

Altogether, our findings reveal that complex multicellularity can emerge and diversify through the interplay of deeply conserved cellular processes with rapidly evolving gene repertoires and regulatory innovation, providing a broader framework for understanding independent origins of organismal complexity.

## Methods

### Biological material and conditions

To cover the breadth of brown algal diversity (**Fig 1a, Fig S1**), as well as to provide a transcriptomic foundation for functional research in brown algae, we focused our species selection on representatives of distinct phylogenetic groups, in particular, emerging model organisms belonging to speciose clades (such as Laminariales, Ectocarpales and Fucales). In addition, we selected species that capture unique biology, such as the rapid change in sexual systems within the genera *Desmarestia, Fucus* and *Saccorhiza* ^20^. With these considerations, we selected the following species for the brown algal expression Atlas: *Ectocarpus* species 7, *Scytosiphon promiscuus, Hapterophycus canaliculatus, Sphaerotrichia firma, Fucus serratus, Fucus distichus, Laminaria digitata, Saccharina latissima, Desmarestia herbacea, Desmarestia dudresnayi, Saccorhiza polyschides, Saccorhiza dermatodea, Sphacelaria rigidula* and *Dictyota dichotoma*.

To standardise naming across the 14 brown algal species, we categorised multicellular samples by generation: gametophyte (“GA”) and sporophyte (“SP”). We further categorised gametophytes and sporophytes composed of a few cells (< 20) or embryonic stages as “early”, sexually immature gametophytes and sporophytes as immature (“imm”) and sexually mature gametophytes and sporophytes as mature (“mat”). Unicellular stages were categorised into gametes, as well as non-gametic mitospore (produced in plurilocular sporangia) and meiospore (produced in unilocular sporangia). Sex of non-gametic unicellular zoids were labelled based on parent sexual genotype. Meanwhile, for the outgroup *S. ischiensis*, we categorised two main stages, unicellular zoids and simple multicells.

Brown algal material was collected either from wild populations (field) or laboratory cultures and quickly rinsed with filtered and autoclaved seawater. Large and rough tissues (such as holdfast, stipe, meristem, and vegetative and reproductive tips) were dissected into ~0.5-1 cm pieces and transferred to 1.5 mL low-bind Eppendorf tubes. These tubes were snap-frozen in liquid N_2_ and stored at −80 °C until processing. Further information on sampling and/or culturing of brown algal specimen used in this study is provided in **Table S1**.

### RNA extraction and sequencing

#### RNA extraction

To generate the poly(A) mRNA transcriptomic dataset, we first dry-grinded the snap-frozen algae while keeping Eppendorf tubes in liquid N_2_ or dry ice. Next, the tubes were removed from the liquid N_2_ or dry ice and immediately thereafter we added 750 µL of preheated (65°C) RNA extraction buffer, consisting of 100 mM Tris-HCl pH8.0 (Thermo Fisher AM9856); 1.4 M NaCl; 2% Hexade**c**yl**t**rimethyl**a**mmonium**b**romid (CTAB) (Sigma Aldrich, 52365-50G); 20 mM EDTA pH 8.0; 1% β-Mercaptoethanol; 2% Polyvinylpyrrolidon (PVP from plant RNA Isolation Aid (Thermo Fisher AM9690). Samples were then vortexed briefly and incubated at 65°C for 1 minute. If necessary, samples were temporarily placed on ice and reheated at 65°C for 30 seconds prior to further processing, since, at this stage, the tubes can be stored on ice for a few minutes. Next, 250 µL 5M NaCl were added into the tubes and mixed thoroughly. An equal volume of chloroform:isoamylalcohol (24:1) was added, mixed well, and centrifuged at 10,000g for 15 minutes at 4°C. The aqueous phase was removed into RNAse-free tubes and re-extracted with 250 µL pure Ethanol and an equal volume of chloroform:isoamylalcohol (24:1) as before. RNA was precipitated by adding LiCl (Thermo Fisher AM9480) to a final concentration of 4M, together with 1% volume of β-Mercaptoethanol, followed by mixing and incubating at −20°C overnight.

RNA was pelleted by centrifugation at full speed (>18000g) for 45–60 min at 4 °C. The supernatant was discarded, and the pellet (containing the RNA) was washed once with freshly prepared 70% cold ethanol and centrifuged at full speed (>18000g) for 10 minutes at 4°C. The pellet was air-dried for 3-5 minutes and resuspended in 30 µL RNase-free H_2_O. RNA concentration and purity was evaluated using a NanoDrop spectrophotometer, to measure the A260/280 and A260/230 ratios. Optionally, the RNA was further checked by electrophoretic analysis of 100 ng on an 1% 0.5x TAE agarose gel. Residual DNA was eliminated using the TURBO DNase Kit (Thermo Fisher, AM1907) according to the manufacturer’s instructions. Final RNA concentration and size distribution were measured using a Qubit fluorometer (RNA BR Assay Kit, Invitrogen Q10210) and an RNA Nano Bioanalyzer (Agilent 5067-1511). Samples with RIN >7 were considered good and used for downstream RNA-seq library preparation. All the RIN values were between 7.8 to 9.2.

#### RNA-seq library preparation

Bulk RNA-seq libraries were prepared using standard commercial kits following the manufacturers’ instructions. Poly(A) selection for mRNA enrichment was performed using the corresponding NEB kit (#E7490) followed by library preparation using the directional RNA library prep kit from NEB (#E7765). Library concentrations were quantified using the Qubit 1× dsDNA HS assay kit (Invitrogen Q33230), and distribution of insert sizes were evaluated using DNA high-sensitive Kit (Aglient 5067-4626). The sequencing was performed in the in-house genome center on a NextSeq2000 instrument with sequencing kit P3-300 (Illumina). The libraries were pooled for sequencing such that for each library we obtained about 30,000,000 reads, corresponding to about 9GB of data.

For samples with limited material, low-input RNA-seq libraries were prepared using the NEB low input RNA-seq library prep kit (#E6420). For this, small tissue fragments (~50-100 cells) or individual sporangia were isolated and placed on glass slides. Excess sea water was removed, and lysis buffer was added directly to the samples. Tissues were mechanically disrupted by applying a coverslip and smashed with forceps by pressing down quickly. The lysis containing RNA was washed off and collected for the library construction. Further information about the RNA-seq, including kit, protocol and person performing the sequencing is given in **Table S1**. The resulting RNA-seq data have been uploaded to NCBI under BioProject PRJNA1458027.

In total, we amassed 3.42 TB of gzip-compressed sequencing data (639 libraries, 41.0 billion reads). 2.17 TB (63.5%) was generated for this study, 0.97 TB (28.3%) was obtained from recent transcriptomic studies produced by the same people and methodologies as this study ^7,58^, and a further 0.28 TB (8.2%) from other sources ^19,74,75^. The sequencing data was processed uniformly to obtain expression levels for each gene across different developmental and sexual contexts.

### Transcript quantification

To quantify the expression level for each gene model in this dataset, we used the nf-core/rnaseq ^76^, which standardises the pre-processing, quantification and QC steps. RNA-seq reads were mapped against the latest genome and annotation versions ^19,20^. To account for the distinct genome biology of brown algae, we set the maximum intron size in the STAR aligner ^77^ to 50,000 bp. To account for technical bias between samples stemming from the non-uniform sequencing fragment coverage (especially in the 3’ of the gene model), random hexamer priming and GC content bias from PCR amplification, we performed quantification using the salmon software ^78^ with `--posBias`, `--seqBias` and `--gcBias`, respectively. This resulted in library-dependent effective gene lengths for each gene, which we used to obtain the effective length-scaled count and length-scaled transcript per million (TPM) values. To remove spurious gene models with low transcription support, we applied a threshold of mean TPM > 2. This resulted in a more uniform total gene content per species of approx. 10.6k to 18.3k (μ 14.9k, σ = 2.1k), compared to the original 14.2k to 42.3k (μ 22.2k, σ = 7.3k). Due to the scarce isoform annotation in several species, we restricted our analysis to the gene-level rather than the isoform-level.

### Orthology inference

To infer orthogroups across the 15 species, with the earliest divergence recently estimated at 450 Ma we used orthoHMM v0.1.0 ^79^, which is suitable for detecting more remote homologues. In cases of genes with multiple isoforms, we selected the longest isoforms as gene model representatives. For cross-species comparison of gene expression, we summed the expression levels of each gene by its orthogroup, effectively treating the orthogroup as a gene and genes as isoforms. This resulted in expression data for 1,455 orthogroups for the 15 species dataset and 4,188 orthogroups for the sporophyte tissue dataset. We used (length-scaled) TPM values for expression levels in order to account for interspecific differences in gene lengths, not captured by counts. The 1:1 orthologues inferred from orthoHMM were also used, in order to ensure that the findings in this work was not biased by the aforementioned grouping strategy.

### Dimension reduction

To visualise transcriptomic variation across RNA-seq libraries in this study, we first transformed the orthogroup TPM values using vst from DESeq2 v1.44.0 ^80^, followed by projecting the data into 2D *t*-SNE space using the R package Rtsne v0.17 (https://github.com/jkrijthe/Rtsne) on the first 20 principal components (PCs). For both the 15 species dataset and the sporophyte tissue dataset, perplexity was set at 30 (default). When visualising species pairs in the sporophyte tissue dataset, perplexity was set at 10 (due to the small sample size). For the 15 species dataset, we quantified the strength of biological signals (developmental stage, sex, species and tissues) by partitioning these sources of variation using a linear mixed model, as implemented in variancePartition v1.34.0 ^28^, on the same vst-transformed orthogroup TPM values. To ensure consistency, we also repeated the analysis on 1:1 orthologues and across alternative RNA-seq transformations including log2, square-root and rlog from DESeq2 v1.44.0 ^80^.

In addition to visualising the transcriptomic space occupied by RNA-seq libraries, we used *t*-SNE to visualise the co-expression space occupied by genes of different evolutionary ages. To achieve this, we also performed *t*-SNE on the first 20 PCs on log2(TPM+1) transformed gene expression data by transposing the expression matrix. Perplexity was set at 100. Seeds were set to 42 for all *t*-SNE plots.

### Gene age inference and transcriptome age index

To explore the evolutionary signals contributing to the transcriptome dynamics at the gene level, we obtained the previously computed estimates for the evolutionary age of each gene ^19,20^ based on a phylostratigraphic procedure ^81^, using GenEra ^65^. In short, GenEra annotated each gene by the deepest taxonomic node in which a homologue can be found in the NCBI NR database using the ‘ultra-sensitive’ mode of the pairwise sequence aligner DIAMOND v2 ^82^. The resulting gene age ranks (‘phyloranks’) ranges from 1 (oldest; cellular organism) to 13 (youngest; genus and species-level) in *Ectocarpus*, though the maximum phylorank differs between species. The maximum values differ due to the difference in the number of taxonomic nodes with available genomic data. We also used the homology detection failure test ^35^ (as implemented in GenEra) to ensure the robustness of any reported macroevolutionary trends by restricting the analysis to a subset of genes where the inferred age cannot stem from homology detection failure.

To detect where evolutionary younger genes are expressed relative to older genes, we used the transcriptome age index (TAI) ^5^, using myTAI v2.3.2 ^83^, which computes the average gene age for a given stage or tissue sample weighed by the expression level of each gene, *i*.*e*.,

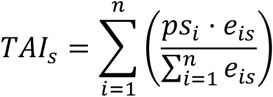

where *ps*_*i*_ denotes the relative gene age (phylostratum) for a given gene *i*. The term *e*_*is*_ denotes the expression level of a given gene *i* at a given sample *s* and *n* denotes the total number of genes. The input TPM values were square-root transformed to balance the need to stabilise the variance of highly expressed genes, while limiting the effect of noise in lowly expressed genes. The statistical significance in the difference of TAI profiles were evaluated via permutation tests implemented in myTAI.

### Expression specificity and breadth

To estimate the expression dynamism of each gene, we extended tau ^33,34^, a commonly used index for tissue specificity, to both tissue and developmental stage specificity for each gene across the entire dataset for each species, *i*.*e*.,

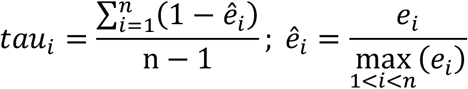

where *n* is the number of samples, *ê*_*i*_ is the expression level of a given gene *i* normalised by the maximal expression value. The input TPM values were summarised using the median across replicates and transformed as log2(TPM+1) to stabilise the variance of highly expressed genes. The resulting tau values range from 0 (broadly expressed) to 1 (tissue or stage specific in expression).

Due to the strong negative correlation between *tau* and mean expression levels, we further visualised the residual *tau* values after accounting for mean log2-transformed expression levels by taking the residuals using `rlm()` (which fits a linear model by robust regression using an M estimator) from the R package MASS v7.3-65. We reported the partial Spearman correlation using `pcor.test()` from the R package ppcor v1.1 ^84^.

### Co-expression network inference and network partition

To explore how the global transcriptome dynamics is structured at the co-expression space, we inferred a consensus co-expression network for each species from 12 different approaches (Pearson correlation, Spearman correlation, PCOR, raw mutual information values, CLR, ARACNE, NARROMI, PLSNET, LLR, SVM, GENIE3, and TIGRESS), using seidr v0.14.2 ^43^, which is inspired by ‘the wisdom of the crowd’ concept in gene network inference ^85^. As input data, we used length-scaled counts from salmon ^78^ which were then transformed via vst from DESeq2 v1.44.0 ^80^, keeping genes with TPM > 2. We used all samples including replicates and excluded species with less than 20 samples. We then used a Bayesian back-boning approach to prune noisy edges in the network ^43,86^. From this consensus co-expression network, we inferred the global position of each gene in the network via the PageRank algorithm as implemented in seidr. As we are interested in relative centrality for cross-species comparisons, we standardised the centrality values as z-scores.

We next partitioned the network into communities (or ‘modules’) to explore the relationship of genes in the co-expression space, using an information-theoretic approach implemented in infomap v2.6.1 ^47^. To ensure consistency between species, we restricted the level used to describe the information flow in the network to two levels via the flag `-2`, along with `--prefer-modular-solution` and `-M 50`. Modules with less than 50 members were excluded from downstream analyses.

To summarise the functional, evolutionary, expression and network profiles of each module, we used information of each gene with regards to gene age, tau, network centrality and inferred protein functions. For protein functional annotation, we used InterProScan v5.74-105.0 ^87^ to infer protein domains and GO terms, as well as publicly available transcription-associated protein (TAP) family predictions ^19,88^. Enrichments of protein domains and GO terms were determined using hypergeometric tests, with the *P*-values adjusted via the Benjamini-Hochberg procedure.

### Cross-species comparison of co-expression modules

To infer the evolutionary conservation of local relationships in the co-expression network, we examined the retention of shared module membership across species, using co-expression modules of the model species *Ectocarpus* as reference. Pairwise orthologues of *Ectocarpus* genes were inferred using reciprocal best hit via the `diamond_rec()` function of the R package orthologr v0.4.2 ^89^ with DIAMOND ‘ultra-sensitive’ mode ^82^. To identify the corresponding modules across species and compute the level of conservation, we used the Jaccard index. This correspondence was visualised across all brown algal species, after excluding cross-species module relationship with Jaccard index < 0.05, using the R package ggsankey v0.0.99999 (github.com/davidsjoberg/ggsankey).

### HGT- and viral-derived genes

Horizontal gene transfers (HGT) represent an additional dimension for genic novelty, alongside taxonomically restricted genes (*i*.*e*. genes without traceable homologues outside of a given taxonomic node – see **Gene age inference and transcriptome age index**). HGT-derived genes in brown algae were obtained from published data, which inferred HGT via homology searches and phylogenetic curation ^19^. In *Ectocarpus*, this dataset is mainly represented by genes derived from bacteria (80/118, 67.8%) and eukaryotes (33/118, 28.0%). The HGT age was then grouped young (Ectocarpus to Ectocarpales), mid (Phaeophyceae to Ochrophyta) and old (Stramenopiles to Chromalveolate) for further analysis.

Given reports of virus-mediated HGTs across eukaryotes ^52^, we further examined genes of putative viral (Nucleocytoviricota; NCLDV) origin characterised by an EsV-1-7 cysteine-rich domain, as proposed in previous works ^51,54^. Since putative viral-derived EsV-1-7 domain genes were not present in the aforementioned HGT list, we first validated the viral affinity of this domain. To this end, we identified EsV-1-7 domain genes (IPR043822) using InterProScan v5.74-105.0 (default parameters) ^87^. Next, we precluded EsV-1-7 domain genes within active NCLDV inserts in brown algal genomes from further analysis due to their presence in the NCLDV insert of *Ectocarpus*. To preclude active NCLDV inserts, we used viralrecall v2.0 (default parameters) to identify potential NCLDV insert regions ^90^. The boundaries of these regions were extended on both sides by 100kb (merging any overlapping regions). Only the enlarged and merged regions with intact NCLDV markers were retained, in order to filter out ‘non-host’ EsV-1-7 domain genes that overlapped with these regions using GenomicRanges v1.56.2 ^91^ and rtracklayer v1.64.0 ^92^.

We then performed pairwise sequence alignment using DIAMOND v2.1.13 ‘ultra-sensitive’ mode ^82^ on ‘host’ genes with EsV-1-7 domains against either the NCBI NR database excluding Ochrophyta (taxonomy ID 2696291), or against a publicly available viral MAG database ^93^. Bacterial and viral sequences were excluded for the NCBI NR search by filtering the ‘sskingdoms’ column for “Eukaryota”. The evalue threshold was initially relaxed for the both searches to 0.1, after which a bitscore threshold of > 40 was applied to mitigate the effect of database size differences. In both searches, the hits with the lowest evalues were retained for each gene. These genes were classified into four categories: those with hits against (1) only NCBI NR Eukaryota (excl. Ochrophyta), (2) only viral MAGs, (3) both NCBI NR Eukaryota (excl. Ochrophyta) and viral MAGs, or (4) none. For genes with both Eukaryota (excl. Ochrophyta) and viral MAGs, the percentage sequence identity (‘pident’) against both databases were evaluated. Having established the sequence homology profile consistent with viral origin of EsV-1-7 domain genes in brown algae, the gene expression patterns of all genes containing at least one EsV-1-7 domain were analysed in terms of their tau and network centrality. Moreover, we computed the proportion of transcripts belonging to EsV-1-7 domain genes in each sample (*TEsV*_*s*_), i.e.,

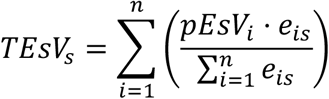

where *pEsV*_*i*_ denotes the presence/absence of EsV-1-7 domain (encoded as 1 or 0, respectively) for a given gene *i*. The term *e*_*is*_ denotes the expression level of a given gene *i* at a given sample *s* and *n* denotes the total number of genes.

In addition, the linear structure of these genes in *Ectocarpus* were visualised with R package gggenes v0.5.1 (https://github.com/wilkox/gggenes) using annotation from Pfam and Coils databases (as obtained from InterProScan), as well as orthogroup information (as obtained from orthoHMM). None of the EsV-1-7 domain containing genes in *Ectocarpus* contained any other domain annotation from Pfam.

### Motif detection

We explored genomic motifs with expression-associated effects by examining the 500 bp upstream of the first CDS of each gene. The CDS was chosen over the transcription start site annotation to maintain consistency in the analysis; 5’ UTR annotation is lacking for most species, as well as a large proportion of genes in the model *Ectocarpus*. The CAGCAG motif (and complementary CTGCTG) was identified using the MEME suite v5.5.7 (using the following parameters: `-dna -mod zoops -minw 3 -maxw 15 -objfun classic -revcomp -markov_order 0`) ^94^. To further validate the significance of the enrichment of the CWGCWG motif, the function `shuffle_sequences()` with euler method from the R package universalmotif v1.22.3 ^55^ was used to re-shuffle the sequences contained within the 500 bp upstream region using three shuffling structure: 1-let, 2-let and 3-let, by adjusting the parameter “k”. The observed number of matches were compared to the empirical null distributions.

To identify potential matches to existing transcription factor binding sites, motif similarity search was performed using TOMTOM v5.5.8 (default parameters) ^94^ against the JASPAR2026 CORE non-redundant database ^95^. Using the full 12-position position weight matrix, no strong matches were identified (lowest q-value = 0.145). Trimming the motif to a 6-position core returned modestly improved similarity (lowest q-value = 0.101), including the three C2H2 zinc finger factor binding sites (MA2532.1, MA1628.2 and MA0697.3).

The cross-species differences of the sequence space in the 500 bp upstream region was assessed by counting 3-let strings for each gene and projecting this count data into a 2D space via PCA using `prcomp()` from the R package stats. The sequence-space PCA and motif enrichment search were repeated with 200 bp data. Genomic data from model eukaryotes were obtained from the ENSEMBL database using biomartr ^96^.

### Motif association with ChIP-seq data

Lastly, we obtained ChIP-seq data from ^58^ and used deepTools v3.5.6 ^97^ to examine the association between the CWGCWG motif with TSS-associated active marks H3K4me3 and H3K9ac marks, as well as a repressive mark H3K79me2. Genome-wide CTGCTG or CAGCAG sites were identified using `vmatchpatterns()` from the R package Biostrings v2.72.1 (https://bioconductor.org/packages/Biostrings), which was then saved as a bedGraph file. These sites were merged and converted to bigwig file format using bedtools v2.31.1 ^98^. Gene annotation in gtf format was converted to bedfile format using `agat_convert_sp_gff2bed.pl` (https://agat.readthedocs.io/). We first computed the positional distribution of CAGCAG or CTGCTG motifs in relation to the gene annotation using the function `computeMatrix scale-regions` from deepTools. We included motif matches 2kb before (`beforeRegionStartLength 2000`) and after the gene body (`afterRegionStartLength 2000`), scaled the gene to 2kb (`regionBodyLength 2000`) to compare the positional distribution across all genes, and excluded regions with only scores of zeros (`skipZeros`). This distribution was visualised using `plotProfile`, using `plotType se` to visualise the standard error). We next computed the positional distribution of H3K4me3, H3K9ac and H3K79me2 coverage with respect to the CAGCAG or CTGCTG motifs using `computeMatrix reference-point`. We centered the reference point (`referencePoint center`), and included ChIP-seq coverage 5kb before (`beforeRegionStartLength 5000`) and after (`afterRegionStartLength 5000`) each CTGCTG or CAGCAG sites. This distribution was visualised using `plotHeatmap`. We used `plotType se` to visualise the standard error around each coverage and set the minimum and maximum values for heatmap intensities using `zMin -3` and `zMax 3`, respectively. To further visualise the overlap between the broad peaks called by MACS2 from the ChIP-seq data (H3K4me3, H3K9ac, H4K20me3, H3K36me3 and H3K79me2), as previously described ^58^, against the genome-wide CTGCTG and CAGCAG sites, we used the function `findOverlaps()` from R package IRanges v2.38.1 ^91^. The proportions of motif sites within and outside each ChIP-seq peak were normalised by the number of nucleotides contained within and outside of each peak, respectively. For gene-level summaries, we used four chromatin signature groups (active, mixed, repressive and null) associated with each gene ^58^, and compared these groupings with the presence/absence of motif matches 500bp upstream.

### Statistical tests and reporting

All statistical tests are reported in the results and figure legends. All permutation tests used to compute the statistical difference in TAI values were done with 50,000 permutations. The R package ecosystems tidyverse v2.0.0 ^99^ (including ggplot2 v4.0.0, dplyr v1.1.4, duckplyr v1.1.1, forcats v1.0.0, tibble v3.3.0, tidyr v1.3.1, stringr v1.5.2, purr v1.1.0) and tidymodels v1.3.0 (https://www.tidymodels.org/) (including corrr v0.4.5 and rsample v1.2.1) were used for data analysis, visualisation and simulation-based statistical inference. All analysis scripts are provided in https://github.com/LotharukpongJS/atlas_brownalga.

## Acknowledgements

We thank Carole Duch*ê*ne, Romy Petroll, Pélagie Ratchinski, Rita Batista, Victor Fernández-Roces, Hajk-Georg Drost and Josué Barrera-Redondo for valuable scientific discussions. We thank Daniela (Lulo) Gallego Calderon for the algal drawings and illustrations. This work was supported by the MPG and the ERC (grant n. 638240 to SMC), the Moore Foundation (GBMF11489) and the Bettencourt-Schuller Foundation (to SMC). JSL thanks IMPRS ‘From molecules to organisms’.

## Contributions

SMC conceived and designed the experiments, supervised the project and acquired funding. MZ, RL, DL, KAB and MH performed experiments, sample collection and the algae culture. FBH and JSL curated the data. ELG and NFP developed the website, with input from FBH and JSL. JSL performed data analysis and visualization with support from FBH and SMC. JSL wrote the original draft with support from SMC. SMC reviewed and edited the manuscript. All authors read and approved the final manuscript.

## Supplementary Tables

**Table S1**. Detail of strains and culture conditions.

**Table S2**. Spearman correlation between gametophytes and sporophytes.

**Table S3**. Gene co-expression network centrality and module.

**Table S4**. Gene co-expression network module annotation in *Ectocarpus*.

**Table S5**. Expression profile of co-expression module 1 in *Ectocarpus*.

## Supplementary figures

**Figure S1.**
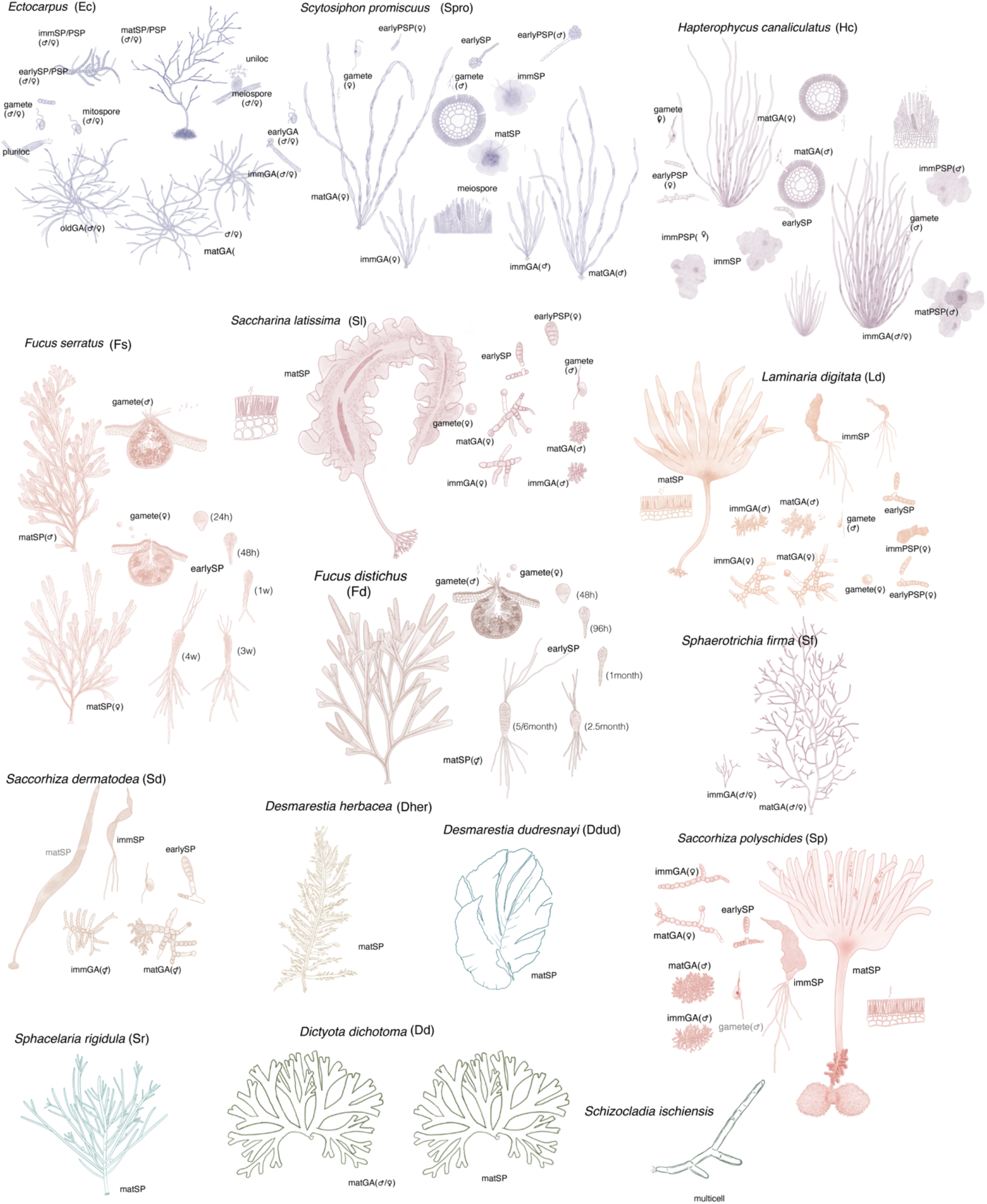
Illustration of species, life cycle stages and tissues in the Seaweed Expression Atlas. Stages not sequenced are labelled in grey. Gametophytes (GA), sporophytes (SP), and parthenosporophytes (PSP) are classified as “early” (<20 cells or embryonic), “imm” (sexually immature) and “mat” (sexually mature). Mitospore and meiospore refer to spore derived from plurilocular and unilocular sporangia, respectively.

**Figure S2.**
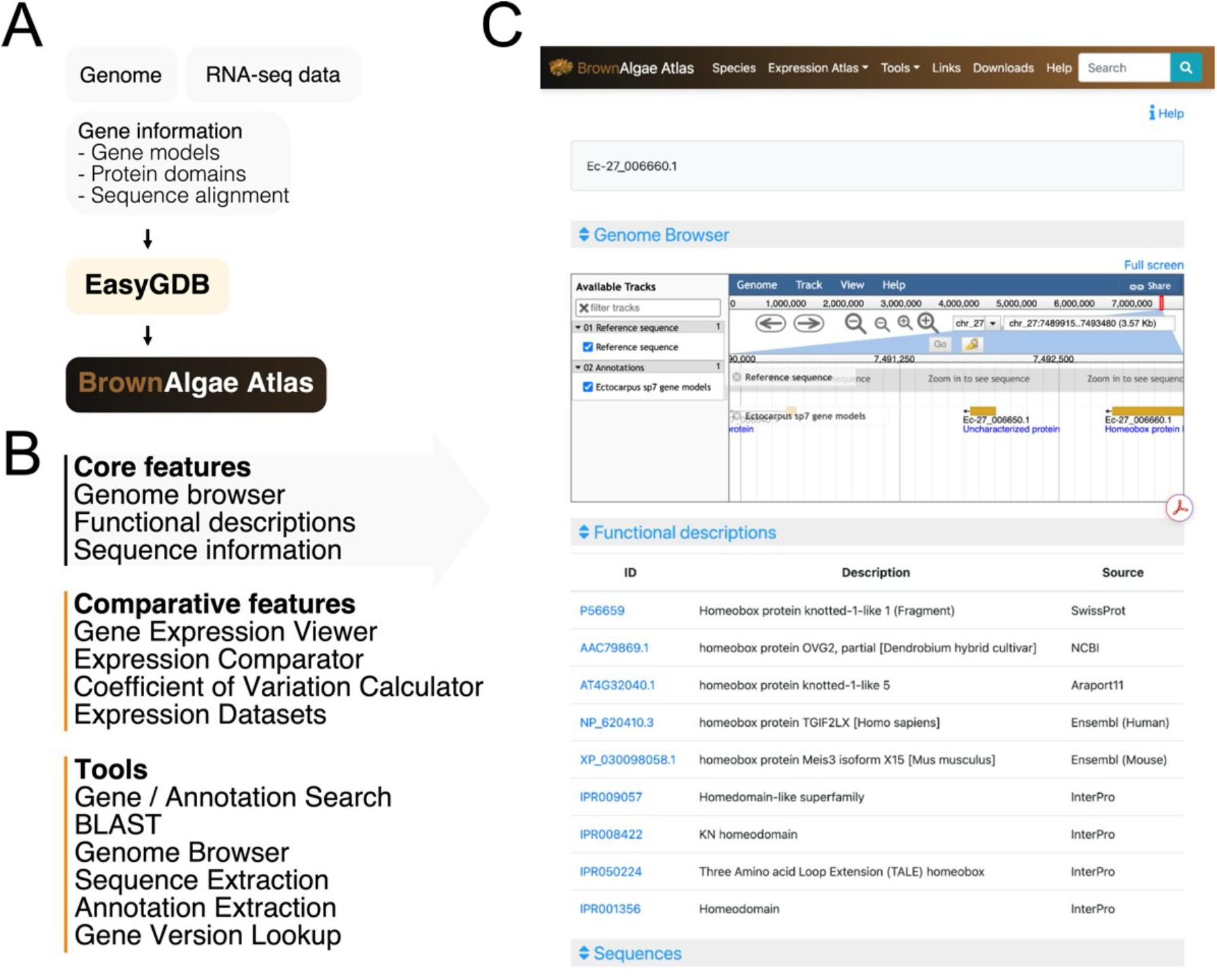
Functionalities of the Seaweed Expression Atlas. (**A**) Workflow to construct the expression atlas website. (**B**) Core and extended features implemented in the Atlas website. (**C**) Gene page example including genome browser and functional descriptions based on sequence alignment matches.

**Figure S3.**
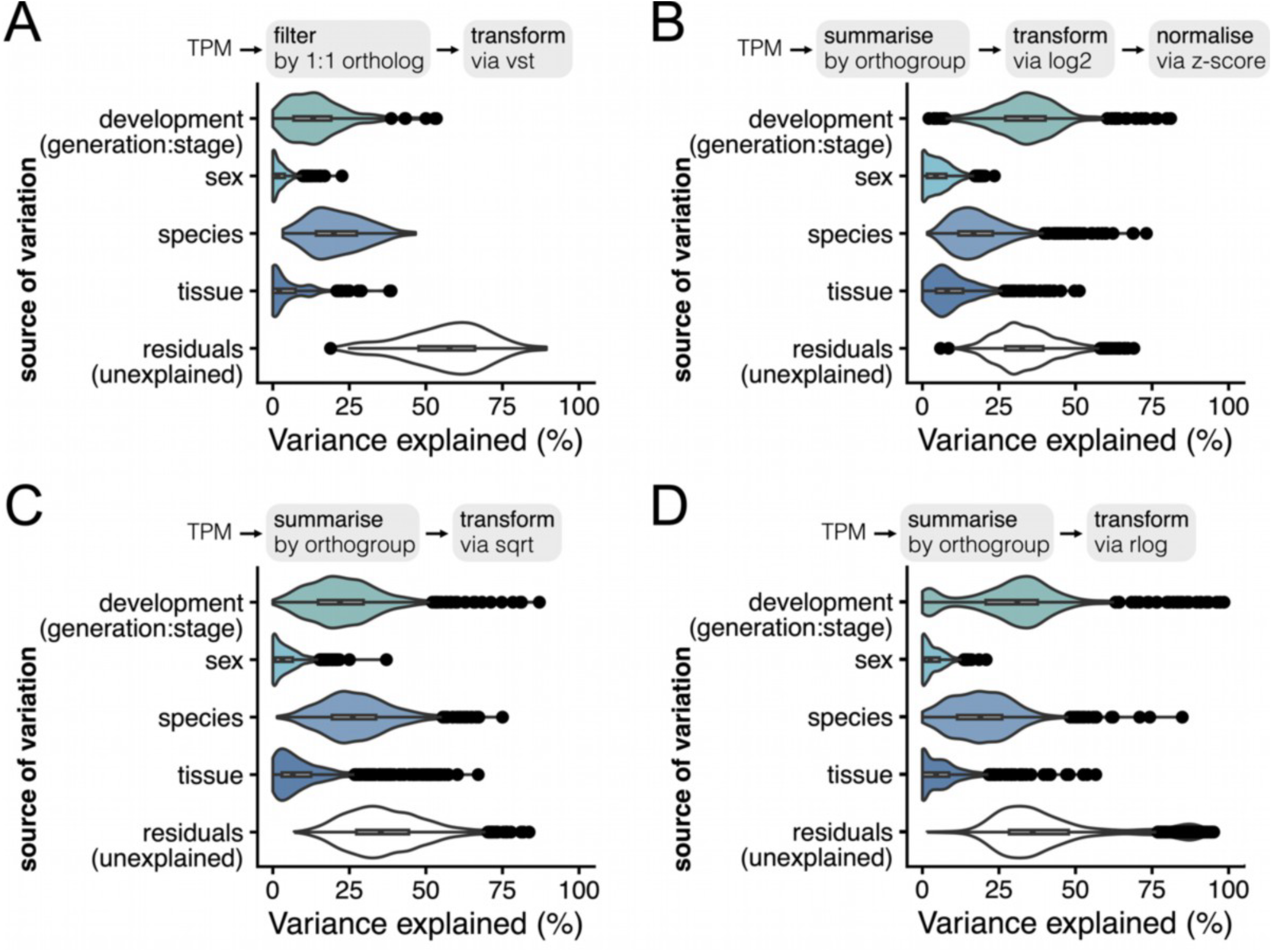
Robustness of variance partitions with regard to orthology information and RNA-seq transformation. Variance partition analysis of (**A**) vst-transformed RNA-seq dataset using 185 1:1 orthologues, (**B**) log2-transformed (*i*.*e*. log2(TPM+1)) RNA-seq dataset using 1455 orthogroups with the resulting transformed value centred and standardised by species, (**C**) sqrt-transformed RNA-seq dataset using 1455 orthogroups, and (**D**) rlog-transformed RNA-seq dataset using 1455 orthogroups. The biological sources of variations considered are development, sex, species and tissue, while the residuals comprise unexplained sources of variation in (**A**-**D**).

**Figure S4.**
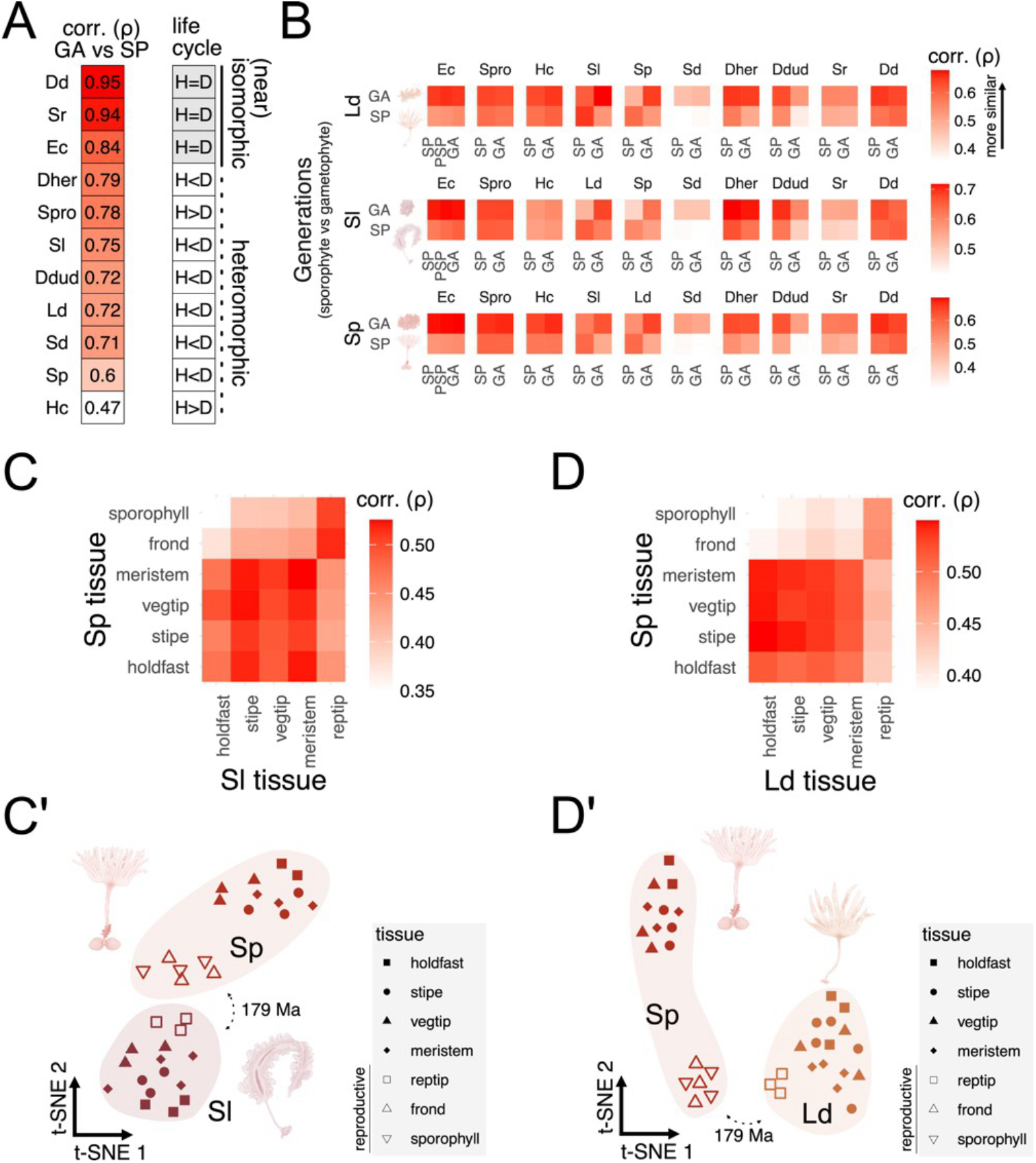
Transcriptome correlation analysis on the gametophyte-sporophyte divergence and convergent evolution of sporophyte forms in Tilopteridales and Laminariales. (**A**) The gametophyte-sporophyte expression correlation within a given species is linked with the dimorphism between the two generations. The average expression profile for each gene within a generation is obtained via median log2(TPM+1). (**B**) Spearman correlation values (ρ) between gametophyte and sporophyte samples across haplodiplontic species *L. digitata, S. latissima* and *S. polyschides*. Higher values indicate stronger correlation, suggesting expression conservation of orthologous genes. GA, SP and PSP denote gametophyte, sporophyte and partheno-sporophyte generations, respectively. For each of the 1455 orthogroups present across all species, the average expression profile at a given generation is obtained via the median vst-transformed expression levels across replicates (after excluding unicells and early developmental stages composed of a few cells). Further comparisons are available in **Table S2**. (**C**-**D**) Spearman correlation values (ρ) between sporophyte tissues alongside a 2D projection (*t*-SNE) (**C’**-**D’**) for (**C**) *S. polyschides* vs *S. latissima*, (**D**) *S. polyschides* vs *L. digitata*. Expression levels across 4188 orthogroups were transformed via vst-transformation for (**C**-**D**). Reproductive tissues are defined as tissues that produce unicells including meiospore in *L. digitata, S. latissima* and *S. polyschides*.

**Figure S5.**
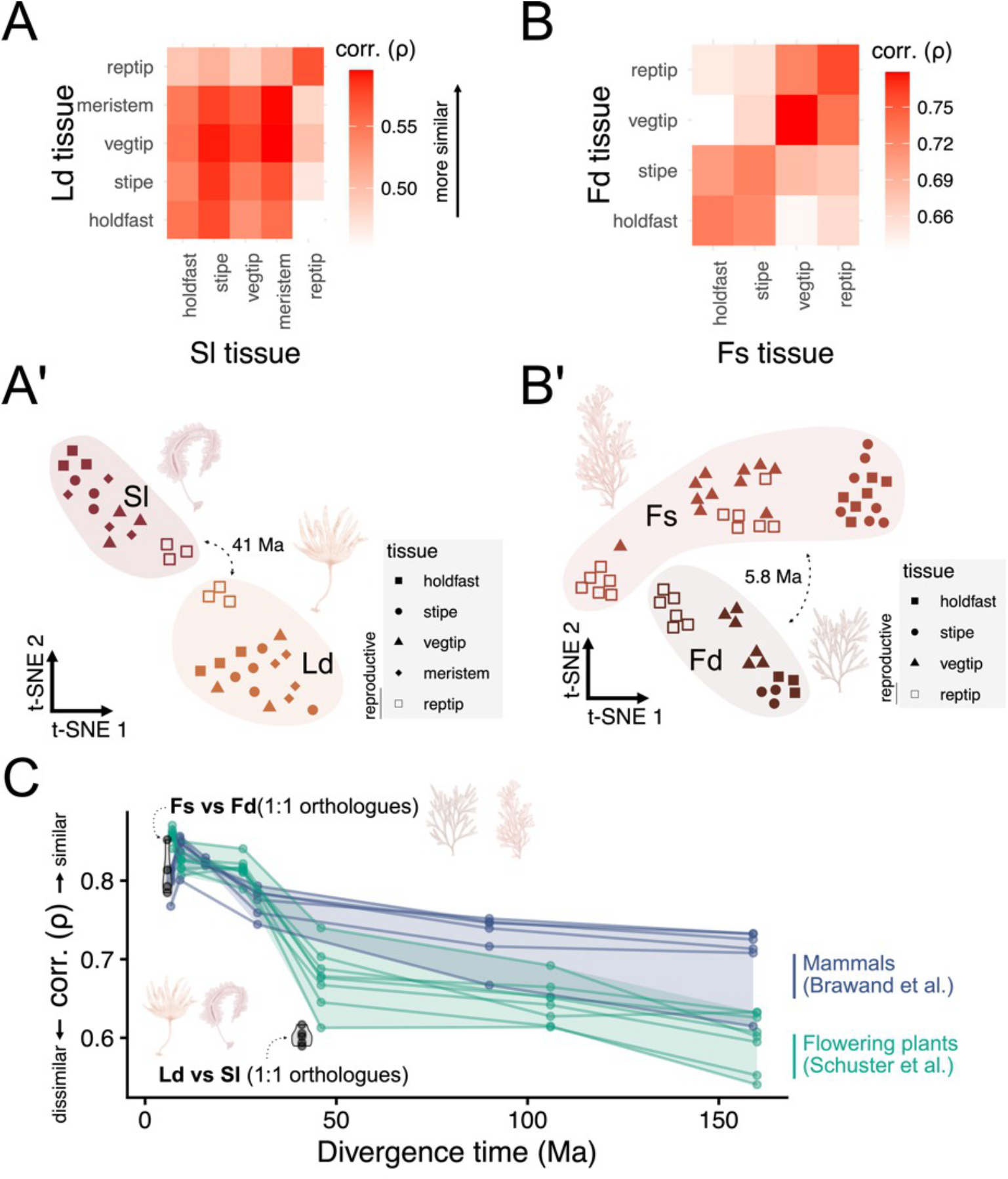
Transcriptome correlation analysis on sporophyte tissue divergence within Laminariales and Fucales. (**A**-**B**) Spearman correlation values (ρ) between sporophyte tissues alongside a 2D projection (*t*-SNE) (**A’**-**B’**) for (**A**) *S. latissima* vs *L. digitata*, and (**B**) *F. serratus* vs *F. distichus*. Expression levels across 4188 orthogroups were transformed via vst-transformation for (**A**-**B**). Reproductive tissues are defined as tissues that produce unicells including meiospores in *L. digitata* and *S. latissima* and gametes in *F. serratus* and *F. distichus*. (**C**) Spearman correlation values (ρ) between the same tissues and/or organs across evolutionary timescales in brown algae (Laminariales and Fucales), land plants and mammals. Inter-species correlations of expression values for 1:1 orthologues in land plant and mammal were obtained from ^32^. The original source of the mammal data is from ^31^, which was reprocessed by ^32^. To ensure uniform methodology, 1:1 orthologues were used for *S. latissima* vs *L. digitata* (n = 4033) and for *F. serratus* vs *F. distichus* (n = 4595).

**Figure S6.**
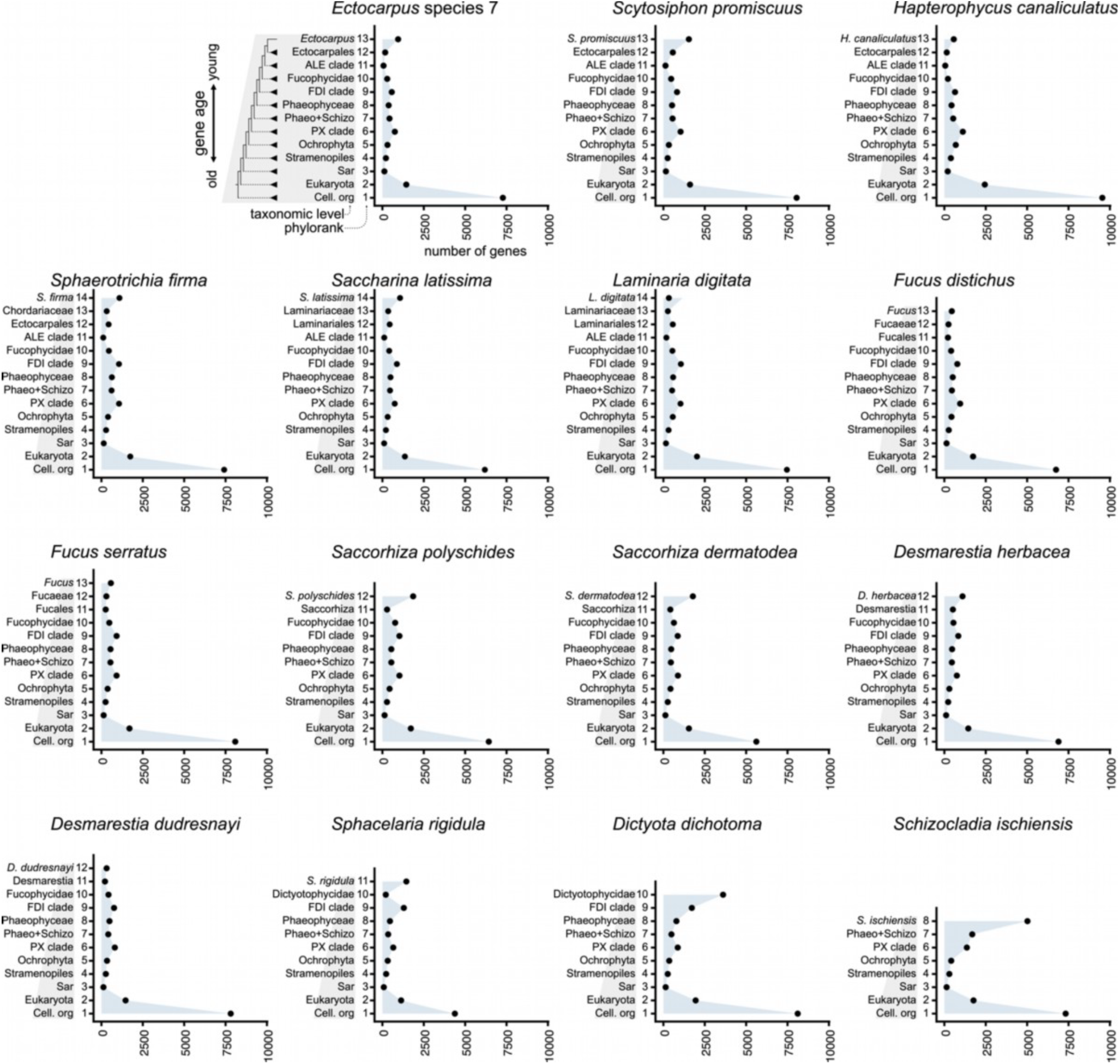
Distribution of genes by evolutionary age (phylorank) across the 14 brown algal species and the outgroup *S. ischiensis*. The x- and y-axes (phylorank and number of genes, respectively) are identical across species and are described in *Ectocarpus* species 7. The taxonomic level “cellular organisms”, which encompass eukaryotes, bacteria and archaea, is shortened to as “Cell. org”. In cases where phyloranks were collapsed, the older corresponding taxonomic node is shown. The maximum phylorank differs among species due to variation in available taxonomic resolution and genomic data.

**Figure S7.**
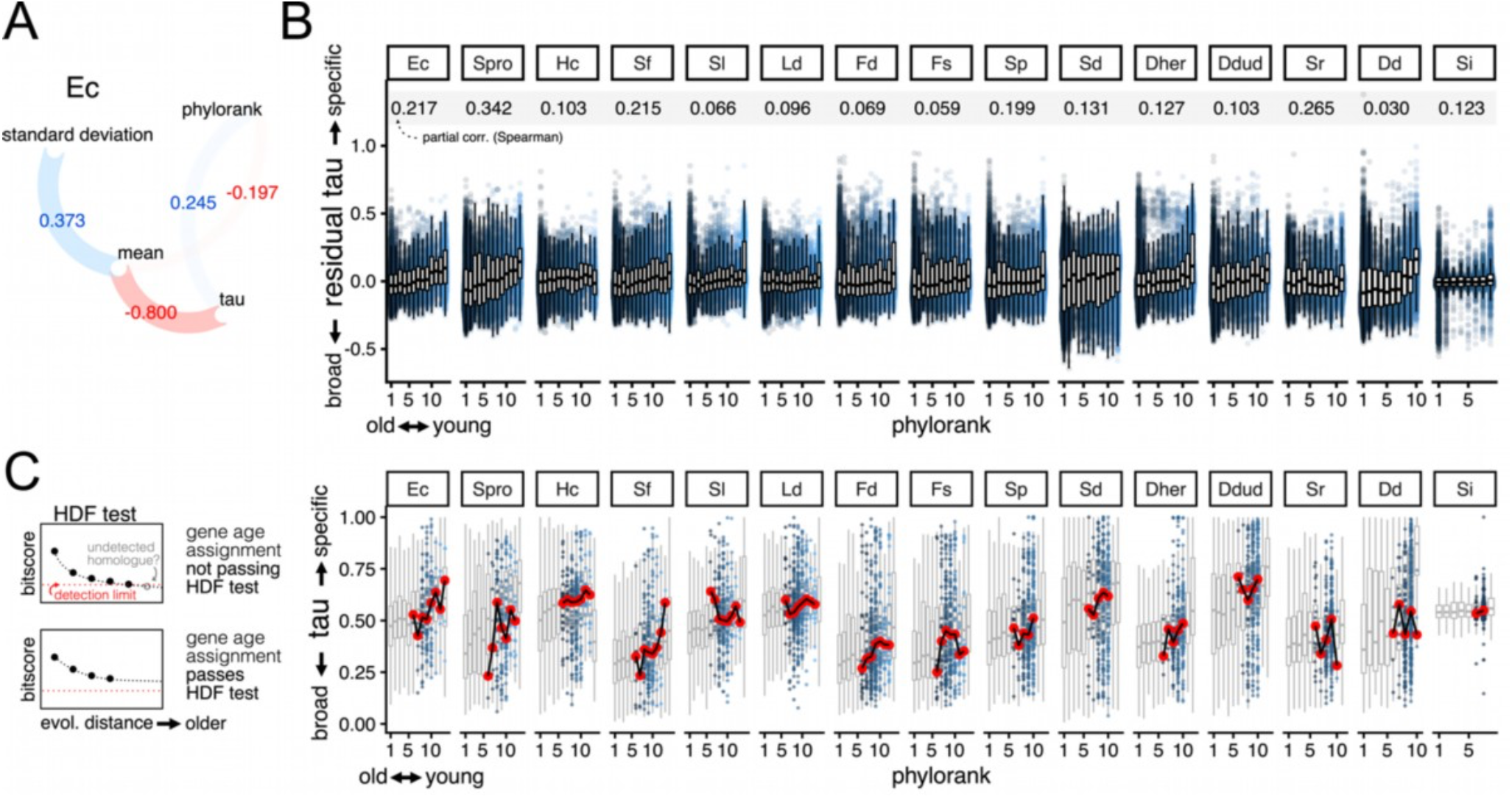
Transcriptomic incorporation of young genes via expression specificity, after accounting for the potential biases in expression specificity and gene age assignment. (**A**) Spearman correlation values between gene age (phylorank), expression specificity (*tau*), mean expression values and standard deviation of expression values. Notably, phylorank is correlated with tau and anticorrelated with mean. (**B**) The relationship between expression specificity (*tau*) and gene age (phylorank) after controlling for the correlation between mean expression and expression specificity. Partial Spearman correlation coefficient is indicated. (**C**) The relationship between expression specificity (*tau*) and gene age (phylorank) on a subset of genes that pass the homology detection failure (HDF) test. HDF test uses the pattern of bitscore decay over evolutionary distance to evaluate the confidence of gene age assignment. The distribution *tau* values by phylorank for all genes is indicated in the grey boxplot. The red points summarise the median *tau* values for genes that pass the HDF test. Genes that pass the HDF test are represented as points. Each point represents a gene, with younger genes coloured by lighter shades of blue in (**B**-**C**). A high *tau* value indicates specific expression (*e*.*g*., stage, sex and/or tissue), while broadly expressed genes are annotated with a low *tau* value. High phylorank indicates evolutionarily young genes, and vice versa for low phylorank.

**Figure S8.**
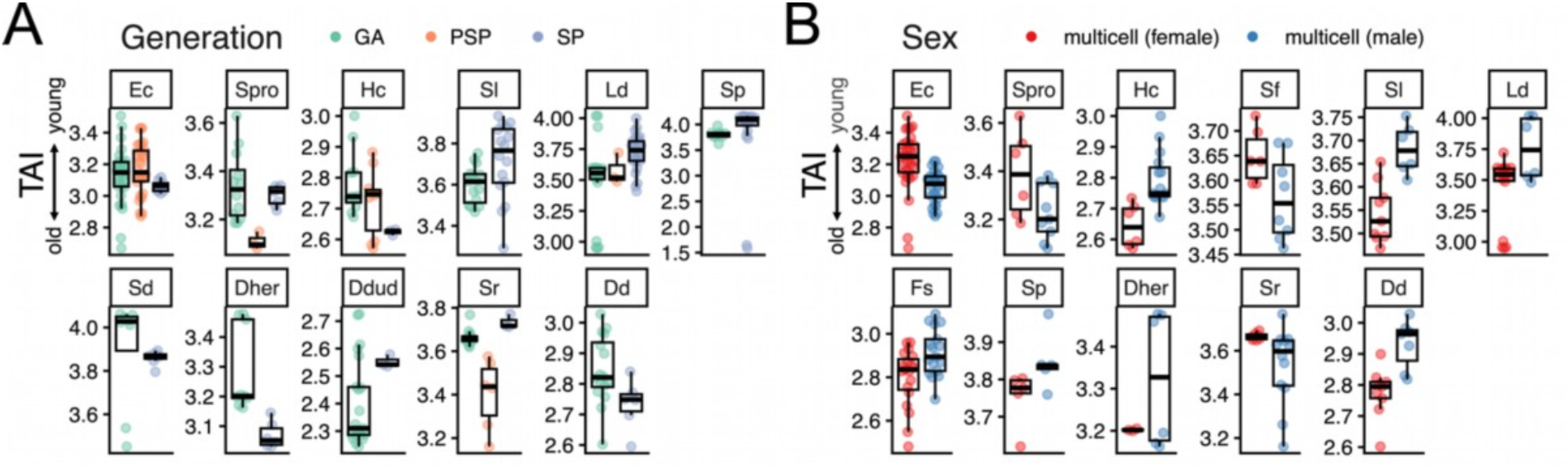
Transcriptome age index (TAI) profiles in multicellular stages. (**A**) TAI profile of multicellular stages grouped by gametophyte vs sporophyte in haplodiplontic species. Parthenosporophyte stages are also included for *Ectocarpus, S. promiscuus, H. canaliculatus, L. digitata* and *S. rigidula*. (**B**) TAI profile of sexual multicellular stages grouped by sex. Non-sexual sporophytes of UV sexual systems are not included. Unicells and early developmental stages composed of a few cells were excluded from (**A**-**B**). TAI values are higher in samples enriched in evolutionarily younger genes compared to those enriched in older genes.

**Figure S9.**
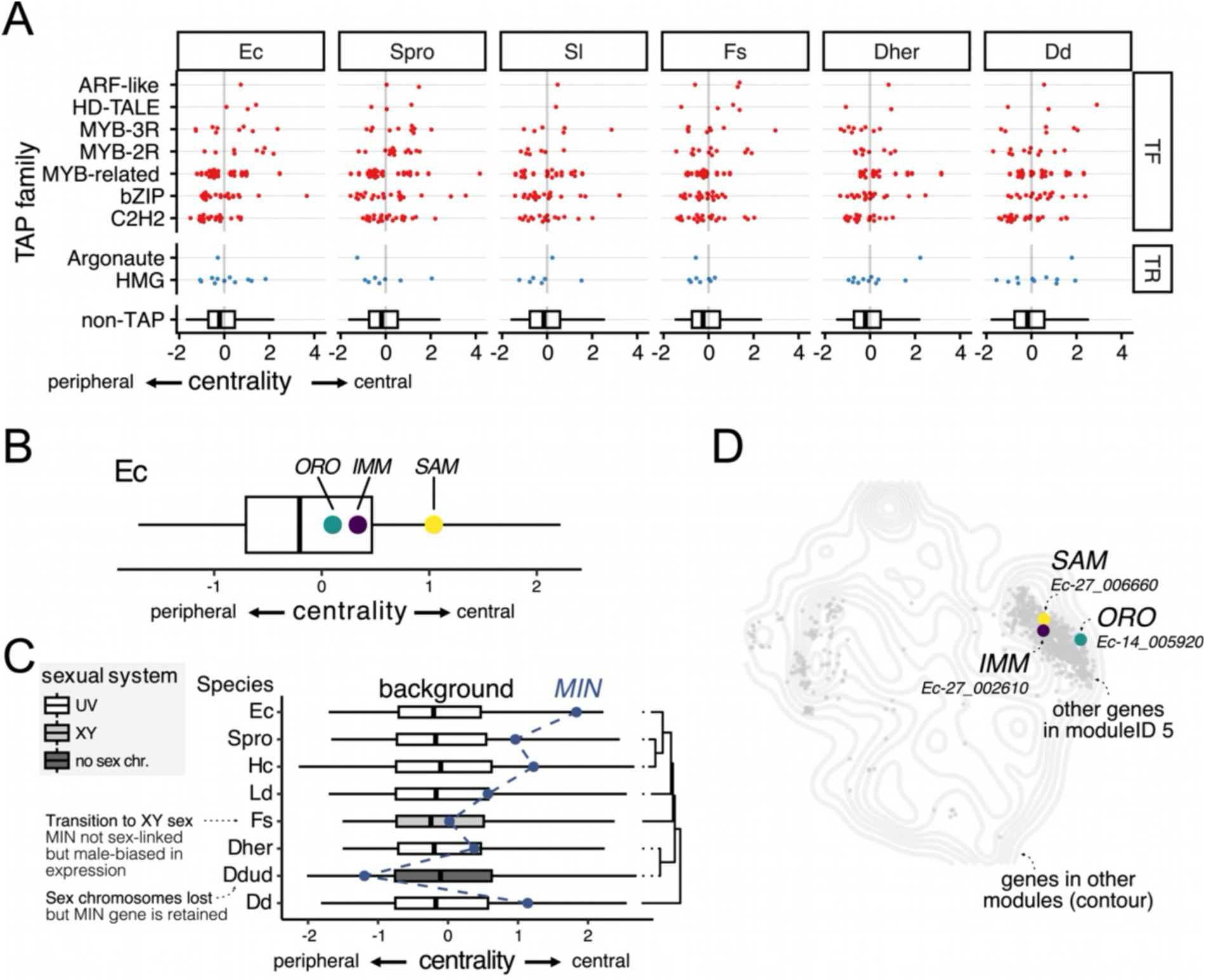
Co-expression network of experimentally characterised genes in *Ectocarpus*. (**A**) Network centrality (z-score PageRank) of a subset of transcription associated protein (TAP) families across brown algal species with associated TAP annotation in ^19^. TAP families are grouped as transcription factors (TF) or transcription regulators (TR). The distribution of other genes in the genome not annotated as TAP is indicated as “non-TAP”. The TAP families are ordered by mean centrality; across the six species, ARF-like and HD-TALE tend to have higher centrality values. (**B**) Network centrality (z-score PageRank) for *Ectocarpus* developmental regulators *ORO, IMM* and *SAM. ORO* and *SAM* contain conserved TALE homeodomains, while *IMM* contains the putatively viral-derived EsV-1-7 domain that is expanded in brown algae. (**C**) Network centrality (z-score PageRank) for orthologous of the *Ectocarpus* male inducer gene (*MIN*) across species with detected orthologues. *MIN* determines male sex in species with UV sex chromosomes. Network centrality of *MIN* gene drops for species that lost the UV sexual system. The evolutionary relationship between the species is displayed in the dendrogram on the right-hand-side. (**D**) A 2D projection (*t*-SNE) of genes in *Ectocarpus* using expression data (as contour), with genes belonging to co-expression module 5 in grey. Module 5 includes *ORO, IMM* and *SAM*, which is labelled in the 2D projection. Each point represents a gene.

**Figure S10.**
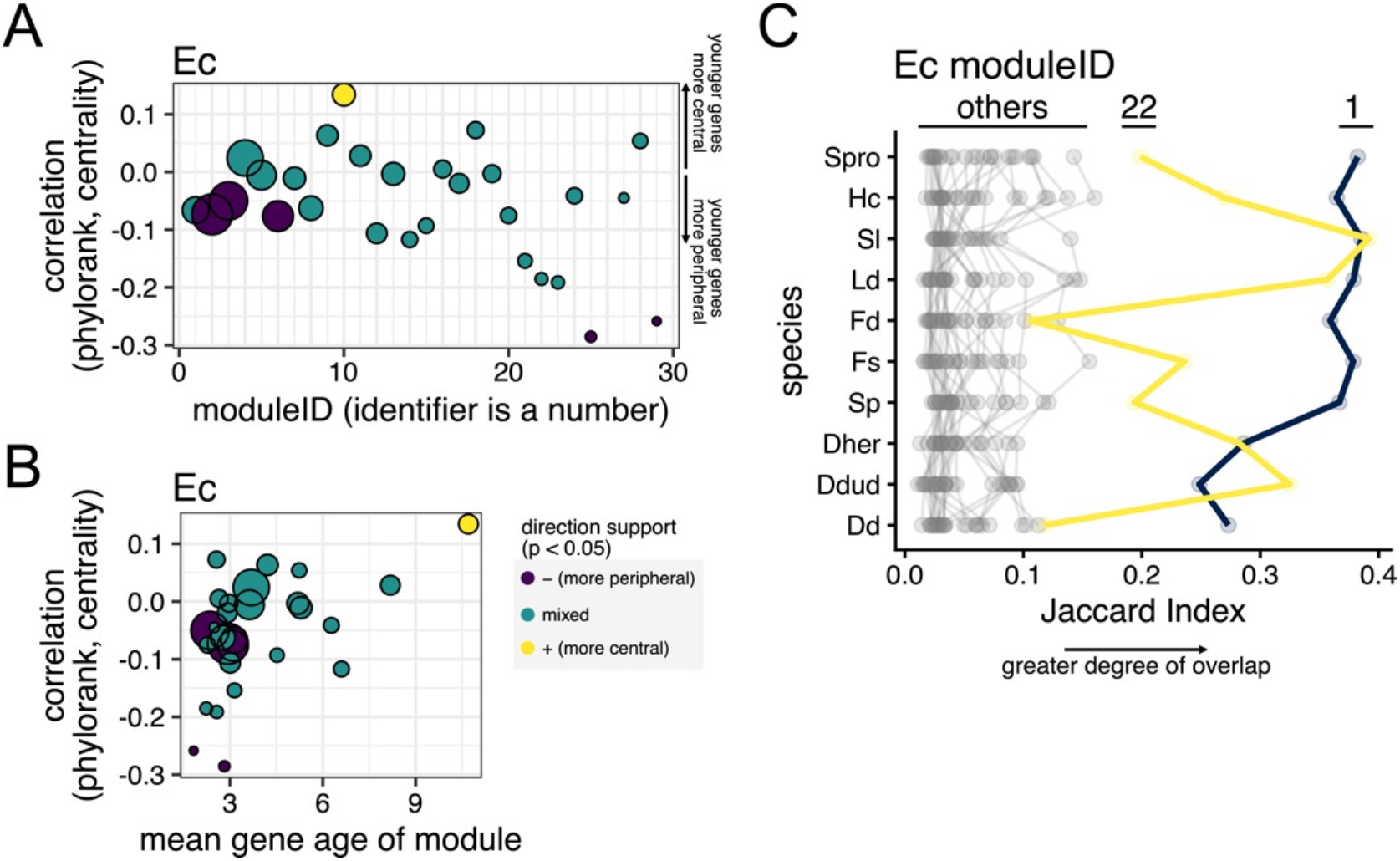
Co-expression network grouped by modules in *Ectocarpus*. (**A**) Spearman correlation between gene age (phylorank) and network centrality (z-score PageRank) across different modules. Module names are given as number. Inconclusive correlations (p ≥ 0.05) are marked as mixed, while significant correlations (p < 0.05) are marked as either positively correlated (younger genes are more central) or negatively correlated (younger genes are more peripheral). (**B**) Spearman correlation between gene age (phylorank) and network centrality (z-score PageRank), by the mean gene age of the module. Modules with older genes tend to have younger genes in the periphery, and vice versa. Mixed, negative and position correlation directions are the same as (**A**). (**C**) Conservation of *Ectocarpus* co-expression modules. Co-expression module conservation was quantified by measuring the overlap of *Ectocarpus* orthologues assigned to the same modules across species using the Jaccard index. A higher Jaccard index indicates a higher module conservation in terms of orthologue overlap. *Ectocarpus* co-expression modules 1 and 22 are have consistently high Jaccard index compared to other modules.

**Figure S11.**
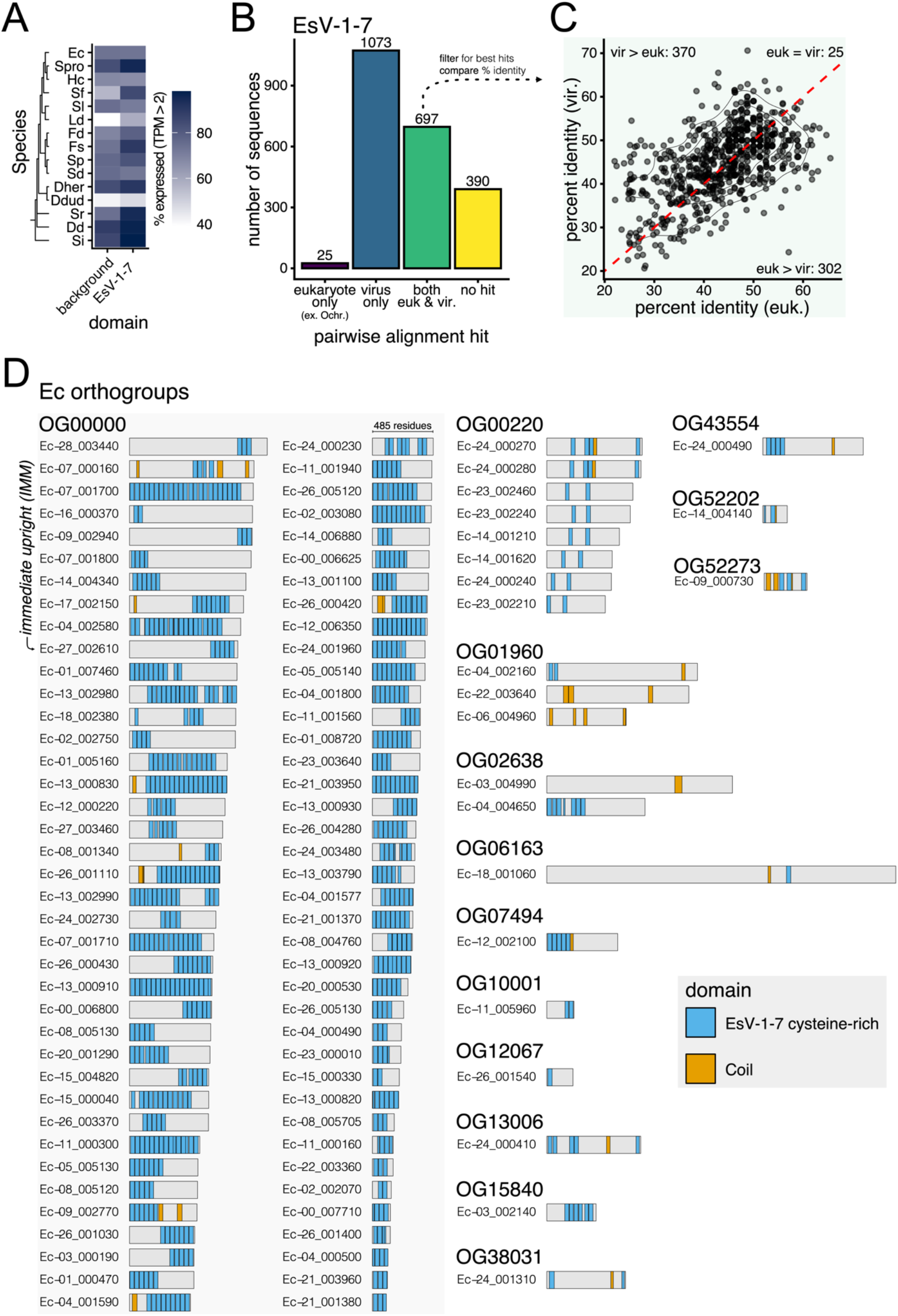
Comparative genomic data on the viral HGT of EsV-1-7 genes and subsequent expansion. (**A**) Proportion of expressed (TPM > 2) genes with EsV-1-7 domain and background across brown algal species. Notably, a higher proportion of EsV-1-7 domain genes are expressed compared to the other genes in the genome in all species except *Ectocarpus, H. canaliculatus, S. dermatodea* and *S. latissima*. (**B**) Number of pairwise sequence alignment hits to eukaryotic and viral protein databases for EsV-1-7 domain genes, with hits grouped by matches to eukaryote (excluding Ochrophyta) only, viral MAGs only, both eukaryote (ex. Ochrophyta) and viral MAGs, and hits to neither databases. (**C**) Difference in percent identity between eukaryotic and viral hits for EsV-1-7 domain genes with matches to both eukaryote (ex. Ochrophyta) and viral MAGs. The density contours represent a bivariate kernel density estimate. (**D**) Gene names within orthogroups that contain EsV-1-7 domains in *Ectocarpus*. Gene lengths are to scale; gene length in amino acid residues is indicated for Ec-24_000230. Genes are annotated from all matches to the Pfam and coil databases. OG00000 is the largest orthogroup across the dataset in this study, which contains the developmental regulator gene *IMM*.

**Figure S12.**
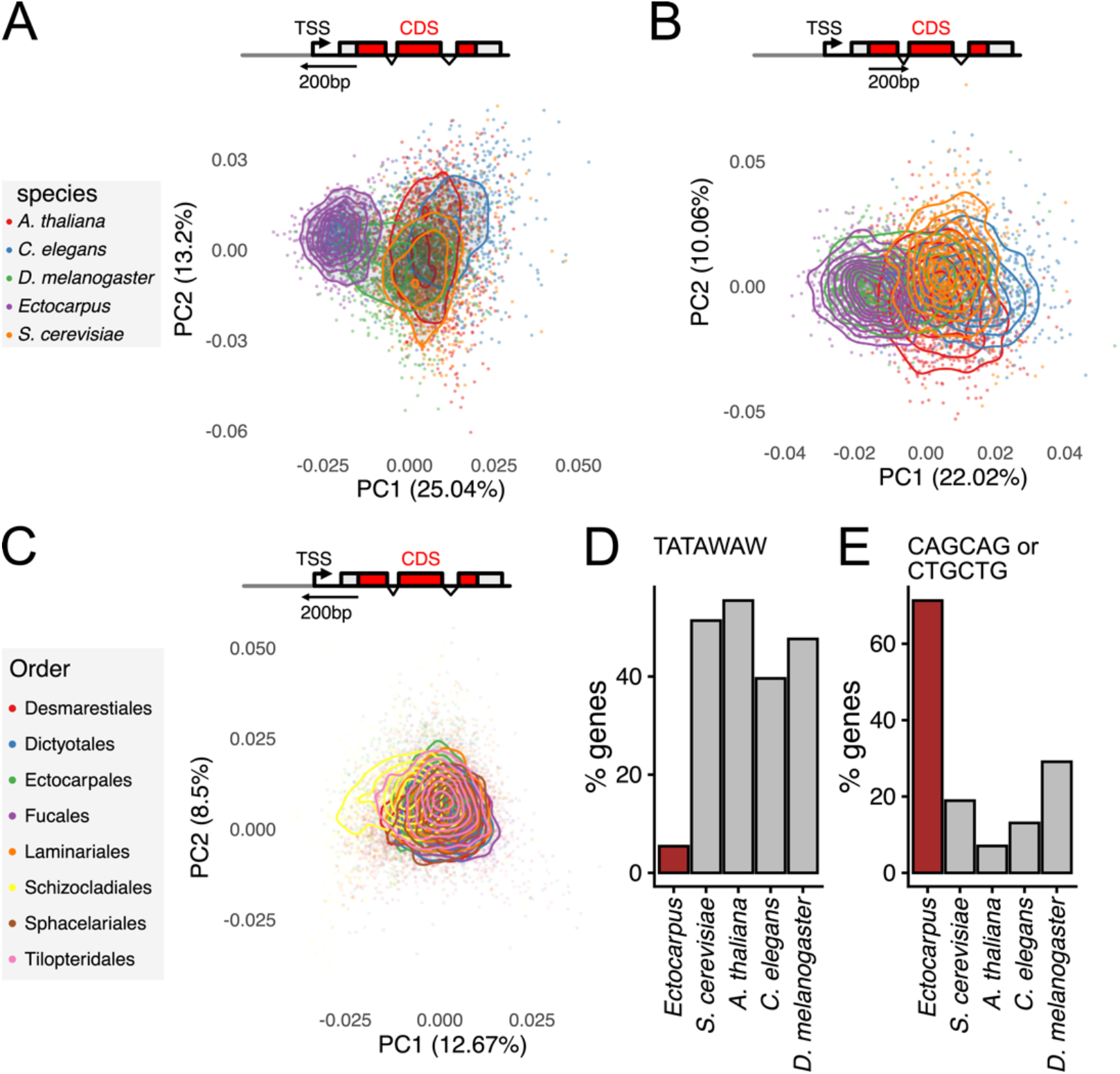
Sequence space of genomic regions proximal to the first CDS across multicellular eukaryotes. (**A**) PCA of 3-let sequence composition 200bp upstream of the CDS across *Ectocarpus* and model eukaryotes (*A. thaliana, C. elegans, D. melanogaster* and *S. cerevisiae*). (**B**) PCA of 3-let sequence composition 200bp downstream of the CDS across *Ectocarpus* and model eukaryotes. (**C**) PCA of 3-let sequence composition 200bp upstream of the CDS across 14 brown algal species and the outgroup *Schizocladia*, coloured by taxonomic Order. Each dot represents a gene in (**A**-**C**). For visualisation, the number of genes displayed has been subsampled to 500 for each species and the density contours represent bivariate kernel density estimates in (**A**-**C**). (**D**) Percentage of genes with exact TATAWAW (W is either A or T) matches 500bp upstream of the CDS across *Ectocarpus* and model eukaryotes. (**E**) Percentage of genes with exact CAGCAG or CTGCTG matches 500bp upstream of the CDS across *Ectocarpus* and model eukaryotes.

**Figure S13.**
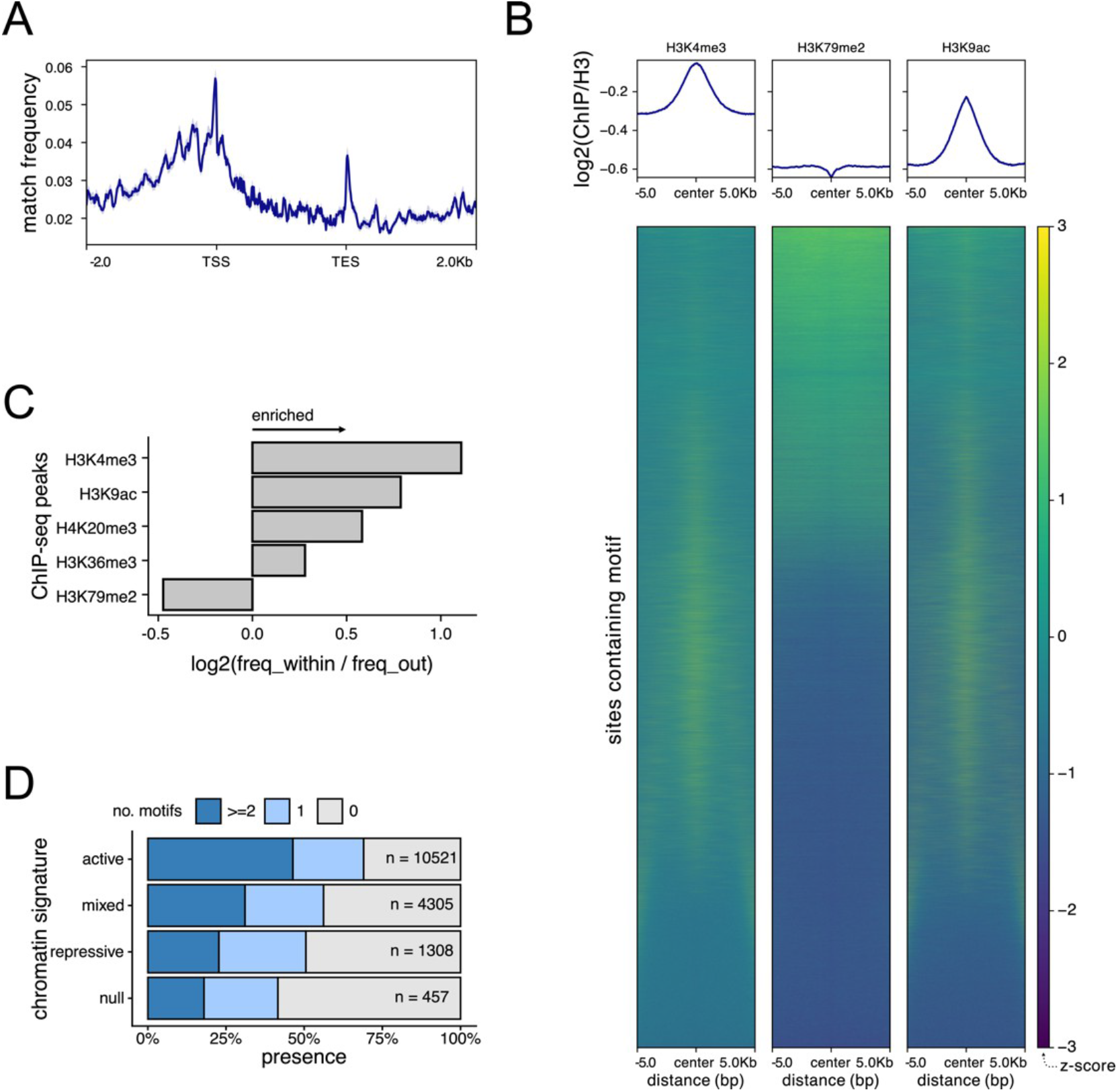
Chromatin and genomic context of CTGCTG and CAGCAG motifs in Ectocarpus. (**A**) Metaplot of CTGCTG or CAGCAG matches over gene models in Ectocarpus. Gene bodies are plotted as proportional lengths, upstream and downstream intergenic regions in kilobases. TSS denotes transcription start site and TES denotes transcription end site. (**B**) Metaplot and heatmap of H3K4me3, H3K79me2 and H3K9ac ChIP-seq coverage surrounding CTGCTG and CAGCAG sites in the *Ectocarpus* genome. Each ChIP-seq signal is normalised by H3 ChIP-seq signal. The position of each motif match is centred within a +/-5kb genomic context. It should be noted that H3K4me3 and H3K9ac are associated with increased transcript abundance and H3K79me2 is associated with decreased transcript abundance at the TSS in brown algae. (**C**) Proportion of CTGCTG or CAGCAG matches within or outside each ChIP-seq peak, normalised by the number of nucleotides contained within and outside of a given ChIP-seq peak. This is done individually for H3K4me3, H3K9ac, H4K20me3, H3K36me3 and H3K79me2. (**D**) Percentage of genes with CAGCAG or CTGCTG matches across genes of different chromatin signatures in Ectocarpus. The total number of genes is displayed for each category. Chromatin data (including H3-normalised ChIP-seq coverage, ChIP-seq peaks and chromatin signature summarisations) were obtained from 58.

